# The mycobacterial selenocysteine machinery: presence and expression

**DOI:** 10.64898/2026.03.03.709296

**Authors:** Phani Rama Krishna Behra, Leif A. Kirsebom

## Abstract

The *Mycobacterium* genus includes more than 190 species that occupies diverse ecological niches. Some are nonpathogenic and environmental, whereas others cause severe diseases both in humans and animals, *e.g*. tuberculosis (TB) and leprosy. SelenoCysteine (SeC) is present in all three domains of life. Here we report the presence of the SeC-machinery (*selA*, *selB*, *selC* and *selD*) genes and selenoprotein (*fdhA*) genes in roughly 40% of 244 mycobacterial genomes.

Their presence is distributed evenly among slow and rapid growing mycobacteria and our data indicate that they were acquired through horizontal gene transfer. Some mycobacteria however lost these genes during the evolution of the genus. We provide RNA-Seq data showing transcript levels of the SeC-machinery genes and *fdhA* in different mycobacteria grown under different conditions. Finally, we suggest that *selC* (the tRNA^SeC^ gene), positioned immediately upstream of *selA*-*selB*, is involved in the regulation of the expression of the SeC-machinery genes *selA-selB*. Together our data expand our understanding of selenocysteine metabolism and its evolution within the *Mycobacterium* genus.

## Introduction

Selenium (Se) was first identified in 1818 by the Swedish chemist, Jöns Jacob Berzelius. For many organisms, selenium is an essential trace element. For humans it is vital and it can influence human health. In 1954 selenium was shown to have a role in microbial metabolism [1,2 and Refs therein]. Among bacteria, selenium exists in three forms: i) as <u>se</u>leno<u>c</u>ysteine, SeC (the 21^st^ amino acid) in specific proteins, which are also present in all kingdoms of life, ii) as a non-covalently bound cofactor and iii) as a modification, 5-methylaminomethyl-2-selenouridine, in certain tRNAs (tRNA^Glu^, tRNA^Lys^ and tRNA^Gln^) [3,4 and Refs therein]. The insertion of SeC into a protein requires an inframe UGA codon, which is generally used as a translational stop codon, and an RNA structural element present in the mRNA, referred to as the <u>se</u>leno<u>c</u>ysteine insertion <u>s</u>equence (SECIS). The size of SECIS is roughly 60 nucleotides and it is folded into a stem loop structure. In bacteria, it is positioned immediately downstream of the UGA codon [5,6]. The UGA codon is read by a specific tRNA, tRNA^SeC^(UCA), and its gene is referred to as *selC* [7]. First, seryl-tRNA-synthetase (*serS*) charges tRNA^SeC^(UCA) with serine generating Ser-tRNA^SeC^(UCA). The serine is then converted to SeC-tRNA^SeC^ by selenocysteine synthase (s*elA*) and selenophosphate (SePO^3-^_3_), which provides selenium. SePO^3-^_3_ is generated from Se and ATP, and this reaction is catalyzed by selenophosphate synthetase (*selD*). The SeC-tRNA^SeC^ binds to a specific elongation factor, SelB (*selB*), which together with GTP, forms a complex that brings SeC-tRNA^SeC^ to the A site on the ribosome that exposes UGA with the SECIS element in its close proximity, which interacts with SelB [8]. The outcome is incorporation of SeC, and proteins having SeC are referred to as selenoproteins. The majority of known selenoproteins are enzymes involved in various cellular functions such as redox reactions [4,9]. In bacteria, SelA, SelB, tRNA^SeC^ (*selC*) and SelD are referred to as the SeC incorporating machinery [3,5,10]. In addition, SelU (gene *selU* or *ybbB*) is involved in mnm^5^Se^2^U synthesis (modification) on certain tRNAs (see above) [9]. Standard annotation systems often fail to identify/predict selenoproteins because UGA is a translational stop codon and among Actinobacteria, to which mycobacteria belong, UGA is the most abundant translational stop codon [11]. Finding selenoproteins can be based on the presence of the SeC-machinery and the presence of SECIS in close proximity to the codon UGA in the mRNA.

Initally, selenoproteins were identified in *Escherichia coli* and in *Clostridium* species by using genetic approaches [3,7,12]. With the advancement of genome sequencing, and the availability of a growing number of bacterial genomes, offer new possibilities to identify the presence of the SeC-machinery and selenoprotein genes [1,13–16]. Comparative genomics has been used to identify the presence of the SeC-machinery and selenoproteins in roughly 20% of sequenced bacterial genomes. Among these, 56 mycobacteria have been reported to have SeC-machinery genes [9,17–25]. We recently reported the presence of the tRNA^SeC^ gene (*selC*) in 102 mycobacterial genomes [26]. Having access to the majority of the mycobacterial genomes as well as several genomes of specific mycobacterial strains [26,27] gave us the incentive to map the presence of the SeC-machinery and selenoprotein genes among mycobacteria. This would provide insight and expand our understanding of the selenocysteine metabolism, and the evolution and distribution of the SeC-machinery within the *Mycobacterium* genus.

We applied a comparative genomics approach using 244 mycobacterial genomes, which included 192 named species representing the *Mycobacterium* genus. We identified genes constituting the SeC-machinery (*selA*, *selB*, *selC* and *selD*) in approximately 40% of the 244 genomes, while no homolog of *selU* could be detected in any of the analysed genomes. Our data further suggest that the SeC-machinery genes have been acquired through horizontal gene transfer. We also provide RNA-Seq data for selected slow (SGM) and rapid (RGM) growing mycobacteria showing the transcript levels for the SeC-machinery genes and the sole selenoprotein gene *fdhA* in response to growth conditions and exposure to different stress conditions.

## Materials and Methods

### Identifying the selenocysteine tRNA^SeC^ and SECIS

To identify the SeC-tRNA in 244 mycobacteria, we used the tools tRNAScanSE [28,29] and Secmarker [22], and the Rfam database [30]. The presence of tRNA^SeC^ genes were searched for using the program tRNAScanSE with default settings. Next, using the specialised tool, Secmarker v0.4, the tRNA^SeC^ gene (*selC*) was identified with the default settings. Also, by using the Rfam database v13.0 along with the infernal; cmsearch command setting the parameter (threshold cutoff) "-T" 45 and run with the default settings identified both "tRNA^SeC^" and "SECIS". Also, SECIS was analysed using the tool bSECISearch tool [31]. Finally, we combined these methods and verified the presence of tRNA^SeC^ and SECIS through manual inspection.

### Selenocysteine, SeC, machinery and selenoproteins

To identify SeC functional protein-coding genes, we used the selenoprofiles v3.6 tool [23], which was designed mainly to identify the selenocysteine machinery in complete and draft genomes. This pipeline consists of three steps, mainly BLAST search, exonerate and genewise and the user can adjust parameters for each profile or all profiles. In our analysis, we set the parameters as per the manual suggested for bacterial genome sequences with default settings and minimum cutoff of 60% identity and query coverage.

All 244 genome sequences were previously annotated using the PROKKA v1.11 annotation pipeline [26] and searched for open reading frames of selenocysteine genes. Data were extracted based on GFF files, with the text search "Seleno or *selA* or *selB* or *selD* or selenocysteine". To confirm missing gene sequences and identify annotation omitted genes, we used BLAST tool blastN search with the cutoff of percentage identity as 70% and query coverage HSP as 60%. We combined these methods and together with manual inspection we identified the presence and absence of SeC-related genes.

### Sequence alignment

The multiple sequence alignments were performed by using the clustalw aligner [32] and the sequences were visualized with the Jalview tool [33].

### Core gene phylogeny

The core gene phylogenetic tree was obtained from Behra et al. [26]. In brief, 387 orthogroups were identified among 244 mycobacterial genome sequences using the SCARAP pipeline. The corresponding protein sequences were concatenated and aligned using the tool MAFFT v7.407 [34]. Phylogenetic tree construction was done using the tool FastTree v2.1.10 [35], by default setting, which uses protein sequence (Jukes canter model). The phylogenetic tree was generated using the tool ITOLtree v3 [36].

### RNA extraction and RNA sequencing

RNA-Seq data (two biological replicates) for selected SGM (four *M. marinum* strains, CCUG20998, 1218R, 1218S and M, and *M. montefiorense* DSM44602) and three RGM, *M. elephantis* DSM44368, *M. mageritense* DSM44476, and *M. monacense* DSM44395/pD5. These mycobacteria were grown in either 7H9 or 7H10 media at recommended temperatures (see Table S3) and total RNA was extracted from exponential and stationary phase cells as reported elsewhere [37,38]. In addition, we analysed our previously reported RNA-Seq data for the *M. marinum* CCUG20998 strain in response to various stressors compared to the control sample [37]. RNA-Seq data for *M. montifiorense*, *M. elephantis*, *M. mageritense* and *M. monacense* grown at different conditions will be published elsewhere.

The respective RNA-Seq short read data were mapped to the respective reference genome using the pipeline Tophat v 2.0.13 [39] and aligned with bowtie2 v2-2-4 [40]. The so obtained aligned BAM files and read counts were generated using the HTSeq v0.9.1 [41] followed by normalization and differential expression analysis using the Deseq2 R package [42], which gives the statistical significance expressed as P+adj values.

To quantify mRNA abundance and compare the expression of different species, we calculated transcripts per million (TPM) values using the TPMCalculator tool [43]. The TPM values were plotted using the R program v3.2.2 [44] in conjunction with the ggplot2 R package [45].

### Visualization of data

Gene synteny plot representing the selenocysteine chromosomal loci in different species were plotted using the genoPlotR v0.8.9 R-package [46]. The structural information of the gene tRNA^SeC^, SECIS elements were plotted using the online tool Rd2t software [47]. The bar plots representing the SeC-machinery and *fdhA* transcript levels correspond to Deseq2 log_2_-fold values and plotted using the R-package ggplot2 [45].

## Results and Discussions

We recently reported the distribution of tRNA and noncoding RNA genes in 244 mycobacterial genomes and we provided data suggesting the presence of *selC*, tRNA^SeC^(TCA), in 102 mycobacterial genomes (Figure 1) [26]. To get further insight into the SeC-machinery and SeC insertion we scanned the mycobacterial genomes to identify all SeC related genes, including the seryl-tRNA synthetase (SerRS) gene (*serS*), required for a functional SeC-machinery. We also searched for the presence of the SECIS structural motif in order to identify protein candidates carrying SeC, refered to as selenoproteins. The presence of homologous genes in mycobacteria was identified using different approaches as discussed below (see also Materials and Methods). Since the focus is on the *Mycobacterium* genus, we name the different mycobacteria as, e.g., *M. marinum* throughout the text.

**Figure 1.**
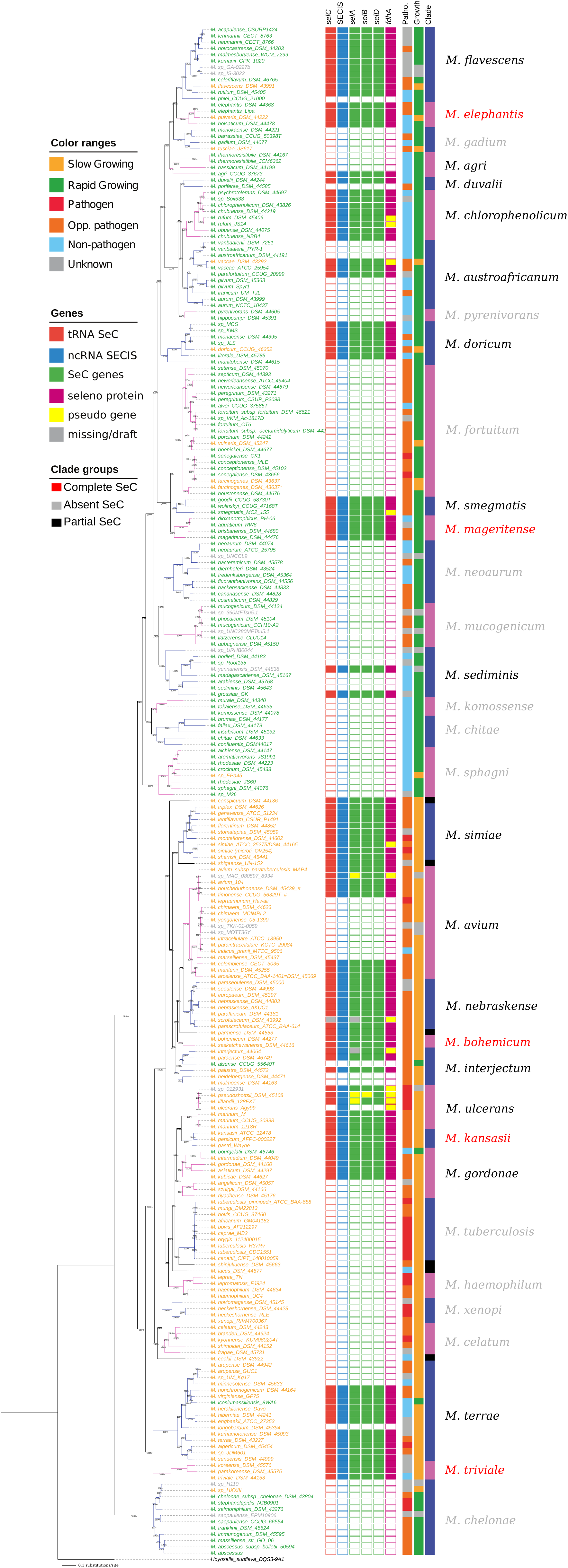
Distribution of selenocysteine genes in mycobacteria Heat maps for 244 mycobacteria showing the presence of selenocyteince machinery genes and selenoproteins based on the combination of predicted results using the tRNAScanSE, Secmarker, prokka, blastN search and selenoprofile search. The representation of mycobacterial core gene phylogenetic tree (387 genes, 244 species) is retrived from Behra et al. [26].

### Presence of homologs and structural characteristics of mycobacterial tRNA^SeC^ (selC)

In addition to the prediction of the presence of *selC* with tRNAScanSE [26,28,29] we searched the 244 mycobacterial genomes with Secmarker [22] and performed a blastN homology search [48]. These approaches resulted in that *M. grossaie* also has *selC*, in addition to the previously identified mycobacteria. Thus, among the 244 genomes 103 (42%) have *selC* (Table S1) including 86 named mycobacteria. The distribution of *selC* among mycobacteria was mapped using the 387-core gene phylogenetic tree (Figure 1) [26]. The data showed that the presence of *selC* is spread evenly between SGM and RGM as we recently reported [26]. Homologs of *selC* were also identified in several other actinobacteria but not in all (data not shown). Interestingly, the *Hoysella subflava* strain DQS3-9A1 [49,50], which is phyologenetically close to the earliest mycobacterial lineage, the *M. chelonae* clade and used as an outgroup in Figure 1, does not have *selC* (data not shown). This is also the case for the members of the *M. chelonae* clade (Figure 1). This indicate that for the mycobacteria having *selC*, the gene was possibly acquired during the evolution of the *Mycobacterium* genus (see below).

The expression of *selC* is predicted to result in an approximately 95 nucleotides long tRNA^SeC^ (Figures 2a and S1a, b). Compared to the common tRNAs, the variable arm of mycobacterial tRNA^SeC^ is longer, which is in keeping with what has been reported for tRNA^SeC^ in other organisms. The "acceptor-T-stem" domains (*i.e*., the combined length of the acceptor- and T-stems measured as the number of base-pairs) in both prokaryotic and eukaryotic tRNA^SeC^ correspond to 13 base-pairs where the acceptor-stem is either 8 or 9 base-pairs long while the number of base-pairs in the T-stem vary between 4 and 5 base-pairs [14,21,22,51,52]. For mycobacteria, the secondary structures suggested that the tRNA^SeC^ acceptor-stem is seven base-pairs long in all mycobacteria while the T-stem is six base-pairs (Figures 2 and S1a-b). Hence, the tRNA^SeC^ "acceptor-T-stem" domain in mycobacteria is 13 base-pairs long consistent with what is known for other organisms.

**Figure 2.**
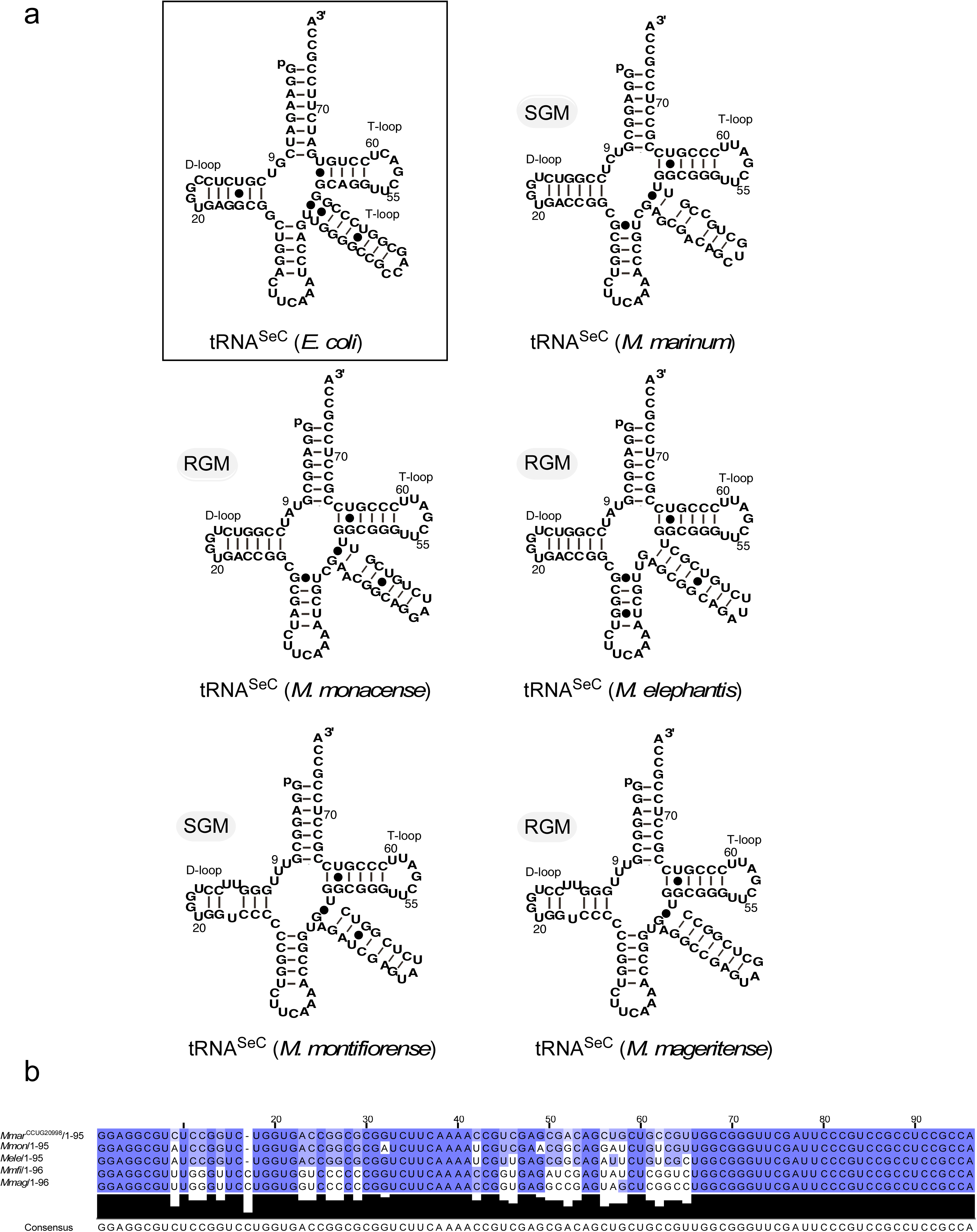
Secondary structures of tRNA^SeC^ and alignment for selected SGM and RGM. (a) Secondary structure of *E. coli* selenocysteine tRNA and predicted secondary tRNA^SeC^ structures of selected SGM (*M. marinum* and *M. montefiorense*) and RGM (*M. monacense, M. elephantis* and *M. mageritense*). (b) Multiple sequence alignment of *selC* for the selected mycobacteria shown in Figure 2a.

For mycobacterial tRNA^SeC^, we also detected structural variations in the junction between the acceptor- and D-stem, in the D-stem and in the variable-arm (Figures. 2 and S1a, b). The D-stem structure with six base-pairs is consistent with available structural information for bacterial tRNA^SeC^ [22,51,52]. However, for some mycobacteria the D-stem structure varies (see Figure S1). Analysis of the mycobacterial tRNA^SeC^ genes further revealed that the identity of the residue positioned immediately 5’ of G_+1_ vary and the majority (60%) carry a G at this position (referred to as N_-1_), while the presence of A, C and U are close to 12, 18 and 10 percent, respectively (Figure S1b and Table S2). Except for *M. conspicium*, *M. gordonae* and *M. grossiae*, all mycobacterial *selC* encodes for the 3’ terminal CCA-motif (Figures 2 and S1b; Table S2). The residue N_-1_, the 3’ CCA-motif and the length of the "acceptor-T-stem" domain play essential roles in the processing of the tRNA 5’ termini by the endoribonuclease RNase P [53, see also Ref 54]. None of the mycobacteria have the tRNA^His^ guanylyltransferase gene, *tgh*, which adds G to tRNA^His^ in eukaryotes after cleavage by RNase P generating an acceptor-stem with 8 base pairs [55; but see Ref 56 for the presence of *tgh* in one *M. phlei* strain]. We also detected sequence variation in the variable loop among the *M. gordonae* clade members (Figure S1b), which might be related to that *selC* was acquired from different sources (see below).

The combined data suggest that structural regions important for mycobacterial tRNA^SeC^ function were identified and these include: i) the amino-stem that interacts with seryl-tRNA-synthetase, SerRS (see introduction and below), ii) the D-loop and D-stem that interact with the N-terminal region of SeC synthase, SelA, and this interaction also discriminates between tRNA^SeC^ and tRNA^Ser^ [7,57,58], and iii) the acceptor- and T-stem that interact with the SeC-tRNA^SeC^ specific elongation factor SelB. Aminoacylation of tRNA^SeC^ is dependent on SerRS (*serS*), which catalyzes the formation of Ser-tRNA^SeC^ that is subsequently converted into SeC-tRNA^SeC^ (see introduction). As recently reported, *serS* is present in all mycobacteria (Table S1) [26].

### Identification of SeC-machinery genes – selA, selB and selD

To convert Ser-tRNA^SeC^ to SeC-tRNA^SeC^ requires two proteins SelA and SelD (see above) [10,59,60]. The presence of the corresponding genes *selA* and *selD* were predicted using three approaches: i) PROKKA annotation followed by extraction of *selA* and *selD* homologs, ii) homologs were identified using blastN searches with *M. marinum* and *M. smegmatis selA* and *selD* sequences as input queries, and iii) selenoprofile search for the SeC machinery and selenoprotein genes based on the Hidden Markov Model, HMM (see also Materials and Methods) [23,48,61].

On the basis of the PROKKA annotation (i), we identified *selA* and *selD* homologs in 94 and 97 mycobacterial genomes, respectively, while the blastN homology search (ii) predicted *selA* and *selD* homologs in 74 and 98 mycobacterial genomes, respectively, and the selenoprofile search (iii) resulted in 91 and 102 mycobacterial genomes having *selA* and *selD* homologs, respectively. Combining the results using these three approaches and by manually verifying the presence of *selA* and *selD* we generated a list of mycobacterial genomes carrying predicted *selA* and *selD* homologs (Figure 1 and Table S1). We used the same work flow to predict the presence of *selB*, which encodes for the SeC-tRNA^SeC^ specific elongation factor. Among mycobacterial genomes, 97, 65 and 97 were predicted to have *selB* homologs using methods i, ii and iii (see above), respectively, and these are listed in Figure 1 and Table S1.

Overall, the presence of the tRNA^SeC^ among the mycobacterial genomes matches the presence of *selA*, *selB* and *selD*. For one mycobacterium, *M. scrofulaceum*^DSM43992^, we predicted the presence of *selD* and *selB* while we were unable to detect *selA* and *selC*, which is likely due to draft genome status. This is probably also the case for *M. interjectum*^DSM44064^, for which we could not identify *selA*. For *M. ulcerans*^Agy99^, we did not detect the presence of any *selA*, *selB*, *selC* and *selD* homologs while the close relative *M. marinum* (four strains; *M. marinum* 1218S not shown in Figure 1) carries these genes. Interestingly, for *M. liflandii*^128FXT^ and *M. pseudoshottsii*^DSM45108^, which are even closer phylogenetically to *M. ulcerans*^Agy99^ [26; note that the *M. ulcerans*^Agy99^ and *M. liflandii*^128FXT^ genomes represent complete genomes (62,63)], we predicted the presence of *selA* and *selB* pseudogenes albeit both carry *selC* (tRNA^SeC^). Data suggest that *M. ulcerans*^Agy99^ diverged from *M. marinum* and the size of its genome has undergone reduction during its evolution [64,65]. The genome sizes for both *M. pseudoshottsii*^DSM45108^ and *M. liflandii*^128FXT^ are also shorter compared to *M. marinum* [26,62,63,66]. This would be consistent with that also these two mycobacteria are subjected to ongoing genome reduction and thereby rationalize the presence of *selA* and *selB* pseudo genes.

The gene synteny shows that in mycobacteria *selC* (tRNA^SeC^) is positioned upstream of *selA* and downstream of *selD* (Figures 3a and S2a). This is in contrast to *E. coli* where *selC* is located elsewhere on the chromosome and distant to *selA* and *selB* [67]. In some mycobacteria, however, a gene with hypothetical function is positioned between *selC* and *selA*. This is the case for *M. flavescens* and *M. elephantis* clade members (Figures 3b and S2a). Moreover, *selC*, *selA* and *selB* are transcribed from the same strand while *selD* is transcribed from the opposite strand (Figures 3 and S2a). For those mycobacteria where *selC* is positioned immediately upstream of *selA*, the distance between the tRNA^SeC^ 3’ CCA sequence and the predicted *selA* translation initiation codon varies and range between 11 and 75 nucleotides (Figure S2b). Within these intergenic regions, we identified Shine-Dalgarno (SD) sequences (*e.g*., 5’-GGAGG) for some mycobacteria. In *M. terrae*, the intergenic region is 11 nucleotides long and the putative SD (5’-AGGAGG) is located one base downstream of the tRNA^SeC^ 3’ CCA sequence. For other mycobacteria, we could not identify any obvious ribosomal binding site (*i.e*., SD) close to the predicted *selA* translation initiation codon, *e.g*., *M. marinum* where the intergenic region is 14 nucleotides long (Figures 3 and S2b). The absence of SD sequence suggests alternative(s) to initiate translation of the *selAB* mRNA in mycobacteria lacking the SD sequence (see also below) [68].

**Figure 3.**
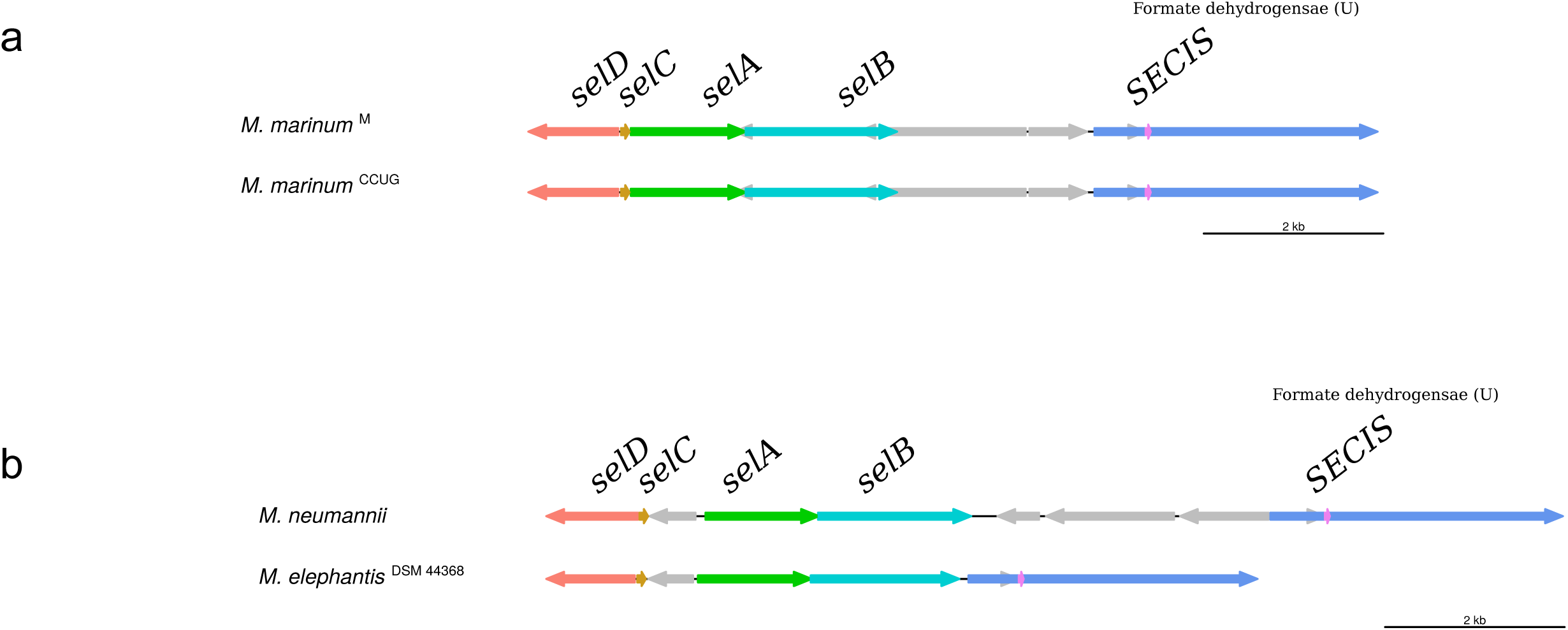
Gene synteny plot showing the selenocystine genes *selABCD*, positioning of the non-coding RNA SECIS element, highlighted in pink, in *fdhA* (the formate dehydrogenase α subunit gene, blue arrow). (a) Two *M. marinum* strains (SGM) and (b) Two RGM, *M. elephantis* and *M. neumannii* as indicated. The colored arrows represent the respective *selABCD* homologs.

### Identification of <u>Se</u>leno<u>c</u>ysteine insertion <u>s</u>equence (SECIS) elements and selenoprotein genes

The selenocysteine insertion sequence, SECIS, is a signature for incorporation of SeC, and SECIS together with the UGA codon are essential to insert selenocysteine into selenoproteins (see above). We used the Rfam database along with the infernal cmsearch tool [14,69,30] and identified SECIS elements in 54 mycobacterial genomes. Moreover, based on a homolog based blastN search [48], we identified SECIS element in 105 genomes (Figure 1 and Table S1).

Following this, we randomly verified some of the SECIS element hits using the online bSECISearch tool [31], which confirmed the predicted sequences as SECIS homologs. The combined Rfam and blastN search data suggested one SECIS element, in *fdhA* (the formate dehydrogenase α subunit gene), per mycobacterial genome. The SECIS element is located immediately 3’ of UGA, at the 3’ end in the corresponding *fdhA* mRNA. Furthermore, searching the 244 genomes using the selenoprofile tool (see above) and blastN homology we predicted the presence of *fdhA* in 92 mycobacterial genomes. Prediction of the secondary SECIS structures for a selected number of mycobacteria revealed similar stem-loop structures with bulges separating the base paired regions (Figure 4; for alignment of mycobacterial SECIS sequences, Figure S3). The mycobacterial SECIS secondary structures show similarities to other bacterial SECIS structures, for example those present in *E. coli fdhF* and *fdnG* mRNAs (Figure 4a) [70]. The *fdhA* UGA is in frame with the annotated initiation codon of *fdoG* (gene for the formate dehydrogenase-O major subunit precursor). Translation (and insertion of SeC) of *fdhA* and *fdoG* transcript/mRNA result in a polypetide with 1064 amino acids (FDH). This is similar to the length of the 1015 amino acids long formate dehydrogenase-O (*fdoG*) in *E. coli* [67].

**Figure 4.**
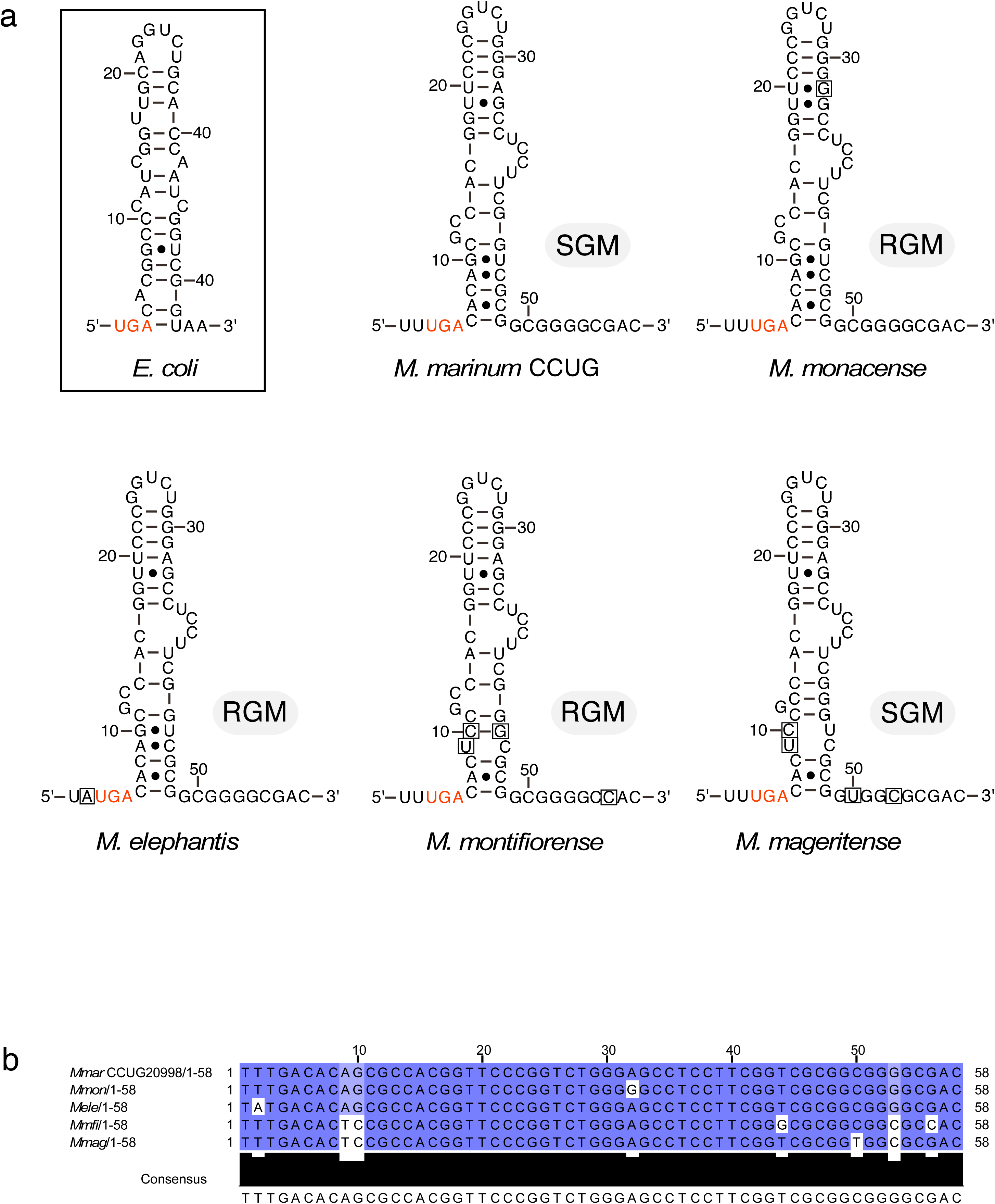
Selenocysteine Insertion Sequence (SECIS) element. (a) predicted secondary structures for *E. coli* and indicated mycobacteria. The required UGA codons are marked in red and box nucleotides refer to differences for the displayed mycobacteria relative to the *M. marinum* SECIS element. (b) Sequence alignment of the SGM and RGM shown in (a).

We conclude that mycobacteria (94 of the 244 mycobacterial genomes; Figure 1) encode for a single full-size selenoprotein, the formate dehydrogenase-O (FDH) α-subunit (note that we identified the SECIS element in 105 mycobacterial genomes, see also below). The FDH is a well-known and dominant selenoprotein homolog in the majority of studied organisms [21]. Its function is to catalyze the reversible oxidation of formate to CO_2_. FDH enzymes can either be dependent on metal ions or metal-independent and the latter are more common among aerobic bacteria [71]. Mycobacteria are aerobic and it remains to be seen whether mycobacterial FDHs indeed are metal-independent. Moreover, noteworthy SelD is a widespread selenoprotein [21], but mycobacterial SelD was not identified as a selenoprotein.

Together, these and the data discussed above suggested that roughly 40% of the 244 mycobacterial genomes contain the SeC-machinery, *selA*, *selB*, *selD*, *selC* (tRNA^SeC^) and SECIS, and one single selenoprotein, FDH (Figure 1 and Table S1). Interestingly, for some mycobacteria we detected the presence of *fdhA* pseudogenes. In particular noteworthy, while *M. ulcerans*^Agy99^, *M. pseudoshottsii*^DSM45108^ and *M. liflandii*^128FXT^ were predicted to have SECIS these mycobacteria carry *fdhA* pseudogenes, likely due to genome reduction during their evolution [26,27,65]. Also, the predicted secondary structure of mycobacterial tRNA^SeC^ have 7 base pairs long acceptor stems, which is in contrast to the 8 base pairs long acceptor stems in other organisms. But, the predicted tRNA^SeC^ secondary structures (Figure 2a) suggested that this is compensated by the six base pairs long T-stems thereby maintaining the 13 base pair long "acceptor-T-stem" domain.

### The mycobacterial SeC machinery and fdhA are likely acquired through horizontal gene transfer

The SeC-machinery, tRNA^SeC^ and FdhA genes were predicted to be absent in several mycobacterial genomes, and among these, in members of the earliest mycobacterial lineage (see above; Figure 1). In addition, gene syntenies suggested that for several mycobacteria carrying these genes, genes with hypothetical functions are located in close proximity of *selC* (Figures 3 and S2a). Together this raises the question whether the SeC-machinery (including tRNA^SeC^) and FDH genes were acquired during the evolution of the *Mycobacterium* genus. For some, however, these genes have been lost after acquisition due to reduction of their genomes. This is exemplified by the analysis of members of the *M. ulcerans* clade (see above; Figure 1). Hence, we generated phylogenetic trees for the individual SeC-machinery and *fdhA* genes to provide further insight and to understand their interrelation.

For *selA* and *selB*, which are transcribed in the same orientation as *selC*, the phylogeny suggested that these genes are grouped in two main clusters, 1 and 2. Cluster 2 can be subdivided into two subclusters, refered to as 2.1 and 2.2 (Figure S4a, b). Mycobacteria belonging to the same clade group together with the exception for members of the *M. gordonae* clade, which group to both clusters 1 and 2. The analysis also revealed that the main clusters (1 and 2) include both slow and rapid growing mycobacteria (SGM and RGM, respectively).

Albeit the short sequence, the phylogeny for *selC* suggested in principle three clusters with two clusters encompassing SGM and one RGM (as for *selA* and *selB*). For *selC*, however, RGM and SGM are separated except for *M. mageritense* clade members (RGM), which are grouped in cluster 1 among SGM (Figure S4c). Also, the *M. gordonae* clade members are separated where *M. gordonae* and *M. asiaticum* belong to cluster 1 and *M. kubicae*, *M. intermedium* and *M. bourgelatii* to cluster 2.

The phylogeny for *selD*, which is transcribed in the opposite direction relative to *selA*, *selB* and *selC* (see above), showed a similar pattern as the *selA* and *selB* phylogeny (cf. Figure S4a-d). For *fdhA*, the phylogeny also suggested separation into two main clusters. But, here cluster 1 and 2 encompassed mainly RGM and SGM, respectively, with a few exceptions (Figure S4e).

These data, together with the gene synteny (Figures 3 and S2a), suggest that the SeC-machinery, tRNA^SeC^ and FdhA genes have been acquired (and lost) during the evolution of the *Mycobacterium* genus.

### Transcription of SeC-machinery genes and fdhA

Having identified the presence of the SeC-machinery genes and the selenoprotein FdhA gene among mycobacteria we were interested in to investigate the transcript levels of these gene (if they are transcribed) in exponential and stationary cells, and in response to exposure to different stress conditions. Hence, we analyzed RNA-Seq data for the *M. marinum* strain CCUG20998 (*Mmar*^CCUG^) grown under different conditions (see Materials and Methods) [37,38]. In this context, we also analyzed the transcription profiles of the same genes for the SGM *M. montifiorense*^DSM44602^ (*Mmfi*) and the three RGM, *M. monacense*^DSM44395^ (*Mmon*^pD5^), *M. elephantis*^DSM44368^ (*Mele*; as discussed above a hypothetical gene is located between *selC* and *selA* in this species) and *M. mageritense*^DSM44476^ (*Mmag*). The RNA-Seq mRNA data for these mycobacteria (*Mmfi*, *Mmon*^pD5^, *Mele* and *Mmag*) were generated using total RNA isolated from exponential growing cells and stationary cells (to be published elsewhere, see Materials and Methods).

The mRNA levels were calculated such that, distribution corresponds to levels expressed as transcription <u>p</u>er kilobase <u>m</u>illion values (TPM) [72] and change corresponds to mRNA levels comparing exponential and stationary growth phase (or changes in response to exposure to different stressors relative to the control; see also [37,38]). The data are shown in Figures 5 and S5. Notable, variation in the transcript levels might either be due to change in expression and/or the result of differences in degradation.

**Figure 5.**
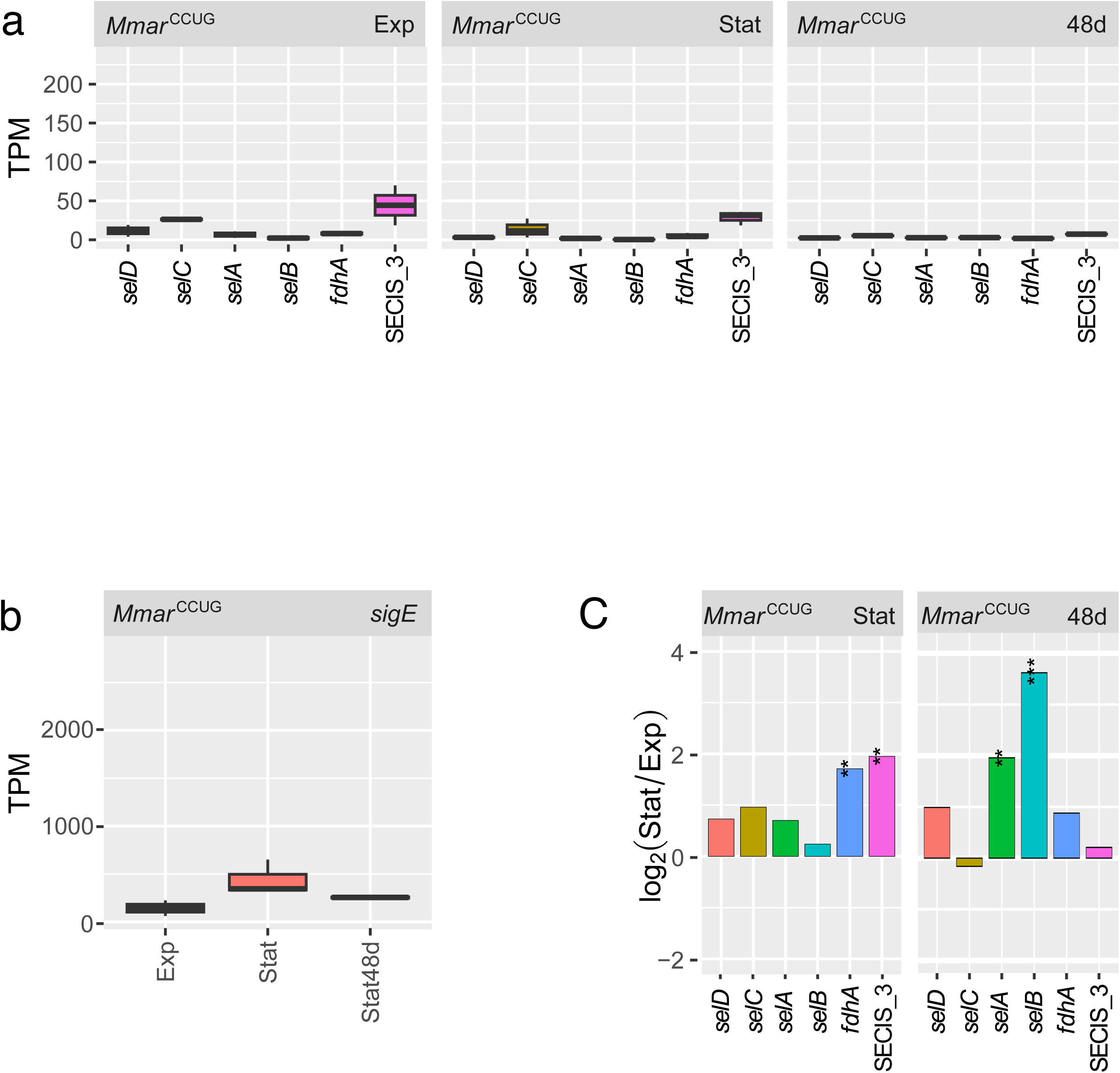
Transcript levels of *M. marinum* CCUG SeC machinery genes and *sigE*. (a) Distribution of SeC machinery, *fdhA* and tRNA^SeC^ transcript levels in exponential, stationary and 48 days old *Mmar*^CCUG^ cells shown as TPM values (see Materials and Methods). (b) Change in SeC machinery and *fdhA* mRNA levels, and tRNA^SeC^ levels, comparing levels in exponentially growing cells with stationary and 48 days old *Mmar*^CCUG^ cells. The values are expressed in log_2_-fold change as indicated. See main text for details and Materials and Methods. Statistical significance: *p+adj< 0.05; **: p+adj < 0.01; ***: p+adj< 0.001. (c) TPM values for *sigE* mRNA levels in exponential, stationary and 48 days old *Mmar*^CCUG^ cells.

For *Mmar*^CCUG^, the SeC-machinery (including tRNA^SeC^) and *fdhA* transcript levels were low relative to *sigE* mRNA. This was observed both in exponential and stationary cells (Figure 5a, c; with respect to mRNA levels for *sigE* and other sigma factors in *Mmar*^CCUG^ [37,38]). When we compared transcript levels in exponential and stationary *Mmar*^CCUG^ cells we detected almost a two-fold (log_2_-fold) increase in *fdhA* mRNA while for the other transcripts the change was small with no change for *selB* mRNA (Figure 5b). However, in late stationary (48 days of growth on 7H10 solid media) *Mmar*^CCUG^ cells we detected larger changes in *selA* and *selB* mRNA levels while no difference was detected for tRNA^SeC^ (*selC*) relative to exponentially growing cells (cf. *Mmar*^CCUG^ Figure 5a, b). For *fdhA* mRNA (48 days), the level was also slightly increased but less increase was detected than comparing exponential and "early" stationary cells. Similar transcript level patterns as for *Mmar*^CCUG^ were also detected in three other *M. marinum* strains [27], *Mmar*^1218R^, *Mmar*^1218S^ and *Mmar*^M^ (Figure S5). Moreover, we also detected increases in SeC machinery mRNA (except for *selD*) and tRNA^SeC^ levels upon exposure of *Mmar*^CCUG^ to starvation (Figure S6). Exposure to other stress conditions such as heat resulted in increased transcript levels of *selA* and tRNA^SeC^ (for other stress conditions see Figure S6 and below).

We conclude that the SeC-machinery and *fdhA* genes are expressed in *M. marinum* with higher levels in stationary cells and under certain stress conditions. Although, the mRNA levels were low in both exponential and stationary cells compared to SigE mRNA levels.

To understand whether the transcript levels is similar in other mycobacteria we studied RNA-Seq data of total RNA isolated from exponential and stationary *Mmfi*, *Mmon*^pD5^, *Mele* and *Mmag* cells (see above). The data are shown in Figure S7. As for *M. marinum*, the TPM values suggested that the transcript levels for *Mmon*^pD5^ (Figure S7a-c), *Mele* (Figure S7d-f), and *Mmfi* and *Mmag* (Figure S7i-h) were low in both exponential and stationary cells (cf. SeC and FDH, and SigE mRNA levels). In contrast to *M. marinum*, the SeC-machinery mRNA and tRNA^SeC^ levels decreased in stationary *Mmon*^pD5^ cells (14 days old cultures) while no or just small changes were detected in 6 and 48 days old *Mmon*^pD5^ cells (Figure S7c). Similar patterns were also observed in 17 and 31 days old *Mele* cells (Figure S7f), while the changes for *Mmag* was at the most modest (Figure S7h). Moreover, transcript levels in exponential *vs*. stationary cells revealed almost no changes in the transcription profiles for the SGM *Mmfi* except for *selD* mRNA, which showed a modest increase in stationary cells (Figure S7h). Together, it appears that upon ageing, exponential *vs*. stationary cells, the transcript levels of the SeC-machinery and *fdhA* genes differ comparing the SGM *M. marinum* (and *Mmfi*) and RGM, in particular *Mmon*^pD5^ and *Mele*.

*Transcript levels of SeC-machinery genes,* fdhA *and tRNA^SeC^ during oxygen depletion* Mycobacteria are aerobic bacteria but they can adapt to survive under limited oxygen supply, *e.g*., during infection in granulomas [73]. An *in vitro M. tuberculosis* model for oxygen depletion, which mimics the situation in granulomas, was developed by Wayne and coworkers [74,75]. We used this model to investigate the SeC-machinery and FdhA mRNA levels as well as levels of tRNA^SeC^ in the SGM *Mmar*^CCUG^ and the RGM *Mmon*^pD5^ in response to oxygen depletion [37 and Ramesh et al., unpublished]. For both mycobacteria, total RNA was extracted at different time points after induction of oxygen depletion and subjected to RNA-Seq (see Materials and Methods). The data are shown in Figure 6a (*Mmar*^CCUG^) and Figure 6b (*Mmon*^pD5^), see also Figure S8a-d.

**Figure 6.**
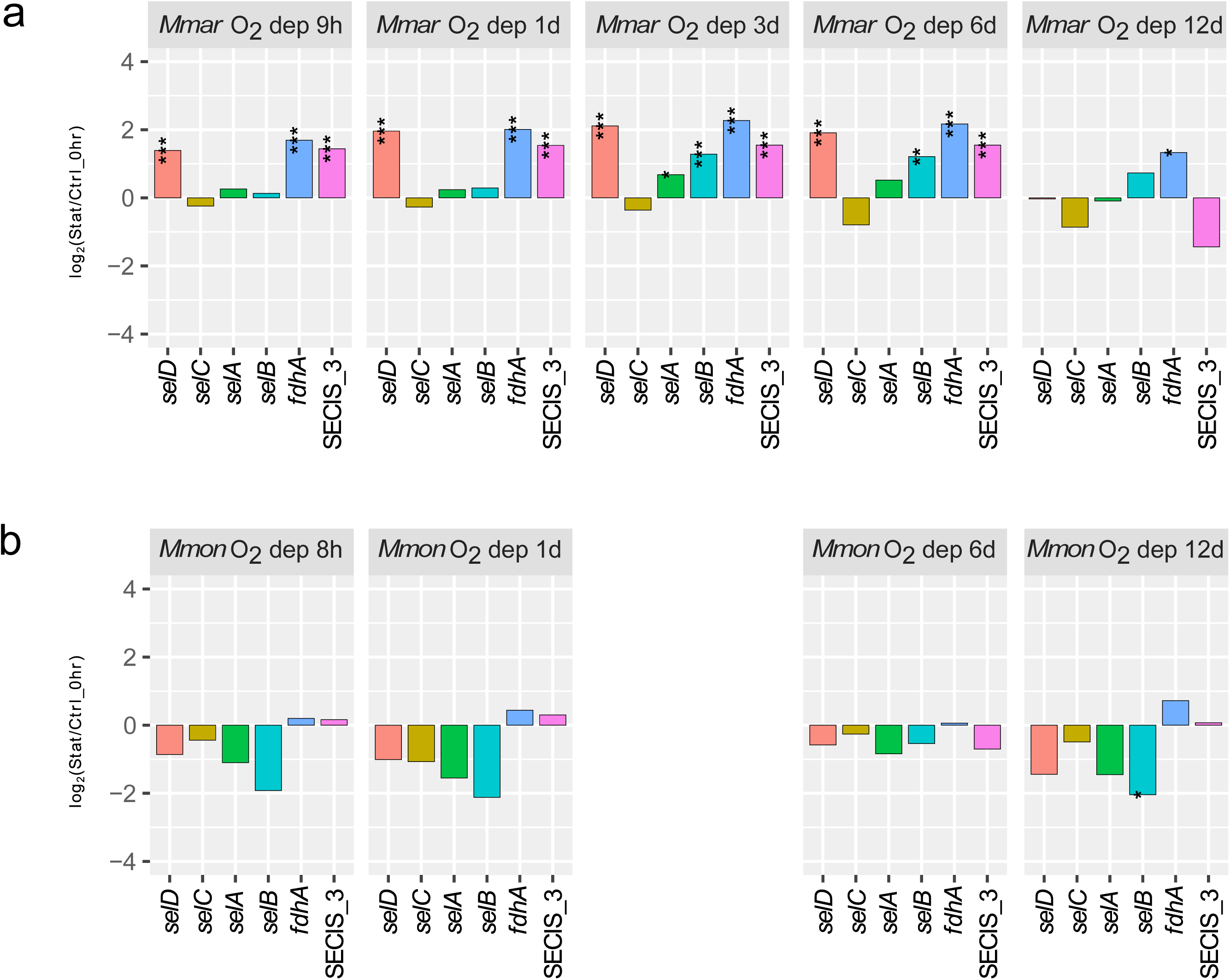
Change in transcript levels of SeC machinery genes and *fdhA* in *Mmar*^CCUG^ and *Mmon*^pD5^ in response to oxygen depletion (micro-aerobic stress). (a) *Mmar*^CCUG^, change in *selA*, *selB*, *selD* and *fdhA* mRNA levels, and tRNA^SeC^ (*selC*) and SECIS_3 levels after 9 hrs, 1 day, 4 days, 6 days and 12 days of oxygen depletion relative to the control, *i.e*. 0 hr. Control corresponds to transcript levels in exponentially growing cells (see Materials and Methods). (b) *Mmon*^pD5^, change in *selA*, *selB*, *selD* and *fdhA* mRNA levels, and tRNA^SeC^ (*selC*) and SECIS_3 levels after after 8 hrs, 1 day, 6 days and 12 days of oxygen depletion relative to the control, *i.e*. 0 hr, see (a). The transcript levels are expressed as log_2_-fold. Statistical significance, see Materials and Methods; *p+adj< 0.05; **p+adj < 0.01; ***: p+adj< 0.001.

For *Mmar*^CCUG^, we detected an increase in *selD* and *fdhA* mRNA at the early time points (9 hours and 1 day), while the level of the *selB* mRNA increased after 3 and 6 days of oxygen depletion. After 12 days, we detected only small variations compared to the control. For tRNA^SeC^, we detected only a small increase over time.

For *Mmon*^pD5^ the response was different. The mRNA levels for the SeC-machinery genes and the tRNA^SeC^ level decreased compared to the control, while no change in the *fdhA* mRNA level was detected. This is similar to the patterns observed in ageing cells (cf. Figure 6b and Figure S7c).

Together, these data suggested that SeC-machinery, *fdhA* and tRNA^SeC^ transcript levels differed in response to oxygen depletion comparing the SGM *M. marinum* and the RGM *M. monacense*.

### Are selC and selAB transcribed together?

For the mycobacteria having the tRNA^SeC^ gene (*selC*) located immediately upstream of *selAB* the distance varies between 11 and 75 nucleotides and for *M. marinum* it is 14 nucleotides (see above; Figures 3 and S2). To understand whether the tRNA^SeC^ gene is transcribed together with *selAB*, paired-end read mapping indicated that this is the case for the four *M. marinum* strains (Figure S9a-d). In contrast, no paired-end readings were detected for *M. elephantis*, which possess a gene between *selC* and *selA* (Figure S9e). For *Mmfi*, *Mmon* and *Mmag* where *selC* is positioned immediately upstream of *selA* (similar to *Mmar*), pair-end read mapping also suggested that *selC* and *selA* are transcribed together (Figure S9f-h).

These data suggest that *selC* and *selAB* are transcribed together in *Mmar*, *Mmfi*, *Mmon* and *Mmag* but not in *Mele*.

### Conclusion and proposed role of having the tRNA^SeC^ gene upstream of selAB

Among the 244 mycobacterial genomes, approximately 40% encode for tRNA^SeC^ (*selC*), the SeC-machinery and FdhA. The genes appear to have been acquired through horizontal gene transfer during the evolution of the genus. For some *M. ulcerans* clade members, however, these genes have been lost due to genome reduction. Furthermore, dependent on mycobacteria the origin of the genes seems to differ.

In *E. coli*, a SECIS-like element is present in the 5’UTR (<u>u</u>ntranslated region) of the *selAB* transcript while *selC* is located elsewhere on the chromosome. The distance between the SECIS-like element and the SelA Shine-Dalagarno (SD) sequence is only three nucleotides (Figure 7a). This element is recognized by SelB and when it is in complex with GTP and SeC-tRNA^SeC^ it binds to the SECIS-like structure thereby represses *selAB* expression [70]. In mycobacteria, no SECIS-like structure could be identified in this region, instead *selC* is present upstream of *selAB*. For several mycobacteria, including *M. marinum*, no apparent SD-sequence is present upstream of the predicted SelA initiation codon while other mycobacteria have a SD-sequence (Figure S2b). In the most extreme case, *M. terrae*, the distance between *selC* and the predicted SelA initiation codon is 11 nucleotides, which includes a SD-sequence beginning one nucleotide downstream of the 3’ CCA end of tRNA^SeC^ (Figure 7b). Our data further suggested that the tRNA^SeC^ gene and *selA* are transcribed together in selected mycobacteria (*Mmar*, *Mmfi*, *Mmon* and *Mmag*; Figure S8). Combined, these findings raise the question whether the tRNA^SeC^ gene or the native unprocessed/processed tRNA^SeC^ transcript is involved in the regulation of *selAB* expression. One possibility is that SelB binds to the native unprocessed tRNA^SeC^ transcript thereby affecting the expression of *selC*, *selA* and *selB*. Compared to *E. coli* this would indicate a similar way to regulate the expression of *selA* and *selB*, except that in mycobacteria this would involve the transcription of *selC* (Figure 7b). Binding of SelB to the unprocessed tRNA^SeC^ transcript might also influence processing and thereby the level of functional tRNA^SeC^. We can also envision that during transcription the GGAGG-sequence in the tRNA^SeC^ acceptor-stem acts as a SD-sequence (consensus 5’-AGAAAGGAGG) interacts with the 30S ribosomal subunit thereby preventing folding and processing of tRNA^SeC^. As a consequence, this would promote interaction with the *selA* SD-sequence (when present). Translational initiation in mycobacteria lacking a *selA* SD-sequence might be achieved through alternative routs [68] and/or involving the GGAGG-sequence in the tRNA^SeC^ acceptor-stem. We also note that that 6C RNA targets, via its C-rich loops, different mRNAs in mycobacteria [76]. Our preliminary data suggest that the SelA mRNA level is affected by the level of 6C RNA in *M. marinum*. It is therefore conceivable that the C-rich 6C RNA binds to the tRNA^SeC^ 5’ G_-1_GGAGGCG-sequence (Figures 2 and S1) during transcription and thereby affect the tRNA^SeC^ and SelASelB mRNA levels. Moreover, in *E. coli* the endoribonuclease RNase P cleaves pre-tRNA transcripts with more than one tRNA in 3’ to 5’ direction [77,78]. We can therefore not exclude that the "*selC-selA-selB*" transcript is first processed by RNase P and other ribonucleases resulting in separation of the tRNA^SeC^ and *selAB* mRNA. Regardless of the way the expression of *selAB* is regulated it is clear from our data that the levels of the SeC-machinery gene transcripts are low and this might be the result of coupling between transcription, tRNA processing and translation. In this context, we note that the folding of the 5’ UTR (operator) of the *E. coli* threonyl-tRNA-synthetase (ThrRS) mRNA is structural similar to tRNA^Thr^. The operator interacts with ThrRS and thereby represses the expression of ThrRS and degradation of the mRNA [79].

**Figure 7.**
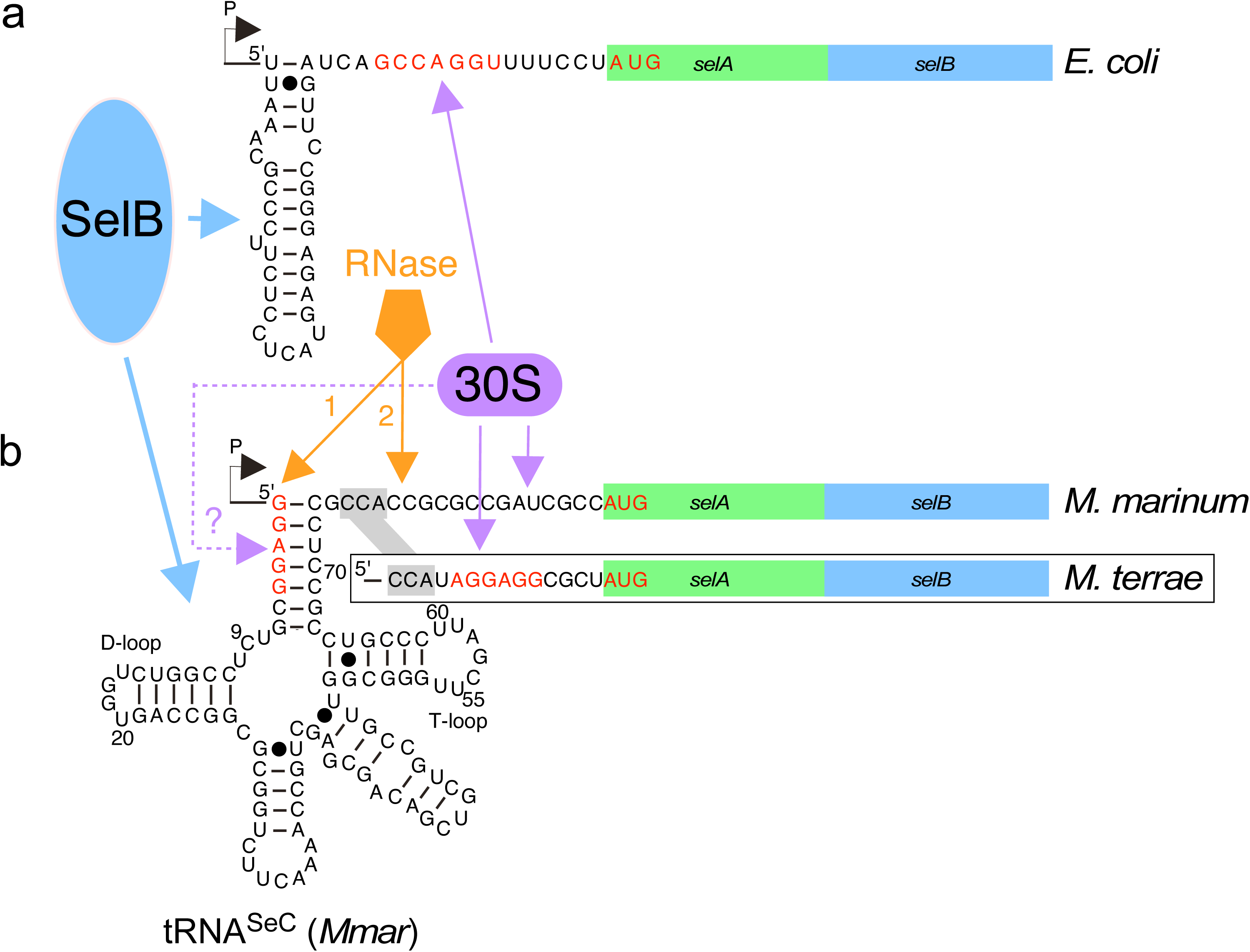
Model of the regulation of *selAB* expression (a) In *E. coli* SelB in complex with SeC-tRNA^SeC^ and GTP (in blue) interacts with the SECIS-like element located upstream of the *selAB* ribosomal binding site (Shine-Dalgarno, SD, sequence marked in red and the ribosomal 30S subunit in purple) resulting in repression of translation of SelAB (see main text and [70]). The black arrow indicates the transcription start, P. (b) The location of the *M. marinum* tRNA^SeC^ upstream of *selAB*, upper sequence, and absence of a ribosomal binding site, SD. The lower boxed sequence refers to the sequence present in *M. terrae*, which includes a ribosomal binding site, SD, marked in red. Residues G_1_ to G_5_ in the acceptor stem are marked in red and the 3’ CCA residues are highlighted in grey. The putative interaction between SelB (in blue) and tRNA^SeC^ is indicated with a blue arrow. The orange marked RNase 1 and 2 arrows correspond to RNases involved in trimming of the tRNA^SeC^ 5’ (RNase P) and 3’ ends. As in (a) the purple 30S ribosomal subunit binds to the SD in *M. terrae* and inititate translation starting at the red marked AUG codon. In *M. marinum* translation initiation of SelAB proceeds by a different mechanism operating on mRNAs lacking SD [68]. Irrespective of binding of the ribosome upstream of the SelAB AUG, the 30S subunit might bind to the tRNA^SeC^ G_1_-G_5_ sequence (which corresponds to a SD sequence) during transcription prior to folding of tRNA^SeC^ (marked by the dashed purple arrow) and thereby affect the translation of SelAB. See also the main text.

To conclude, the way the expression of *selC*, *selA* and *selB* are regulated in mycobacteria differs compare to *E. coli* and it remains to be deciphered. Our present findings establish the groundwork to develop experimental protocols to address this issue.

### Ethics Statement

All methods were carried out in accordance with relevant guidelines and regulations.

## Acknowledgements

We acknowledge our collegues for discussions. Drs B.M.F. Pettersson and M. Ramesh are acknowledged for assistance generating transcriptomic data. Sequencing was performed by the SNP&SEQ Technology Platform in Uppsala and Uppsala Genome Center, which are part of the National Genomics Infrastructure (NGI) Sweden and Science for Life Laboratory. The SNP&SEQ Platform is supported by the Swedish Research Council and the Knut and Alice Wallenberg Foundation. Uppsala Genome Center is supported by the Swedish Council for Research Infrastructures and Uppsala University and is hosted by the Science for Life Laboratory (SciLifeLab). The computations were performed on resources provided by SNIC through Uppsala Multidisciplinary Center for Advanced Computational Science (UPPMAX) under Project uppstore2017058.

## Authors contributions

PRKB and LAK conceived the study. PRKB performed the bioinformatics computations. PRKB and LAK analyzed and interpreted the data. PRKB and LAK wrote the manuscript. The authors read and approved the final version of the manuscript.

## Data availability

The datasets used are available from the corresponding author.

