## Supplementary Figures 1-9 for "The mycobacterial selenocysteine machinery: presence and expression"

Department of Cell and Molecular Biology,
Box 596, Biomedical Centre,
SE-751 24 Uppsala, Sweden

Tel no +46 18 471 4068
Fax no +46 18 53 03 96


*Corresponding author

**Supplementary Table S1a**

Compilation of the presence of selenocysteine machinery and selenoprotein genes 244 among 244 mycobacterial genomes.

**Supplementary Table S1b**

Overall statistical summary of the *Mycobacterium* genus with respect to selenocysteine machinery genes.

**Supplementary Table S2**

Compilation of $N_{-1}$ identity and presence of the 3' CCA in mycobacterial $tRNA^{SeC}$ genes.

**Supplementary Table S3**

Growth conditions for the selected mycobacteria used for RNA-Seq analysis.

**Supplementary Figures S1-S9**

**Figure S1**

(a) Illustration of the *M. marinum* $tRNA^{SeC}$.

(b) Multiple sequence alignment of *selC* (the $tRNA^{SeC}$ gene) present in 103 mycobacteria. The *selC* sequences are grouped according to clades beginning with the *M. flavescens* clade and ending with the *M. triviale* clade, see main Figure 1.

Figure S1a

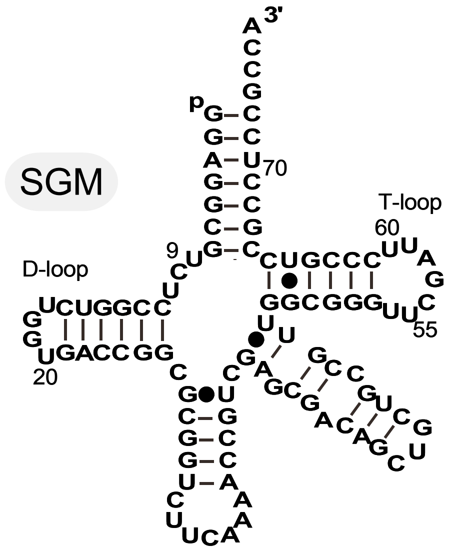


tRNA$^{SeC}$ (*M. marinum*)

### Figure S1b

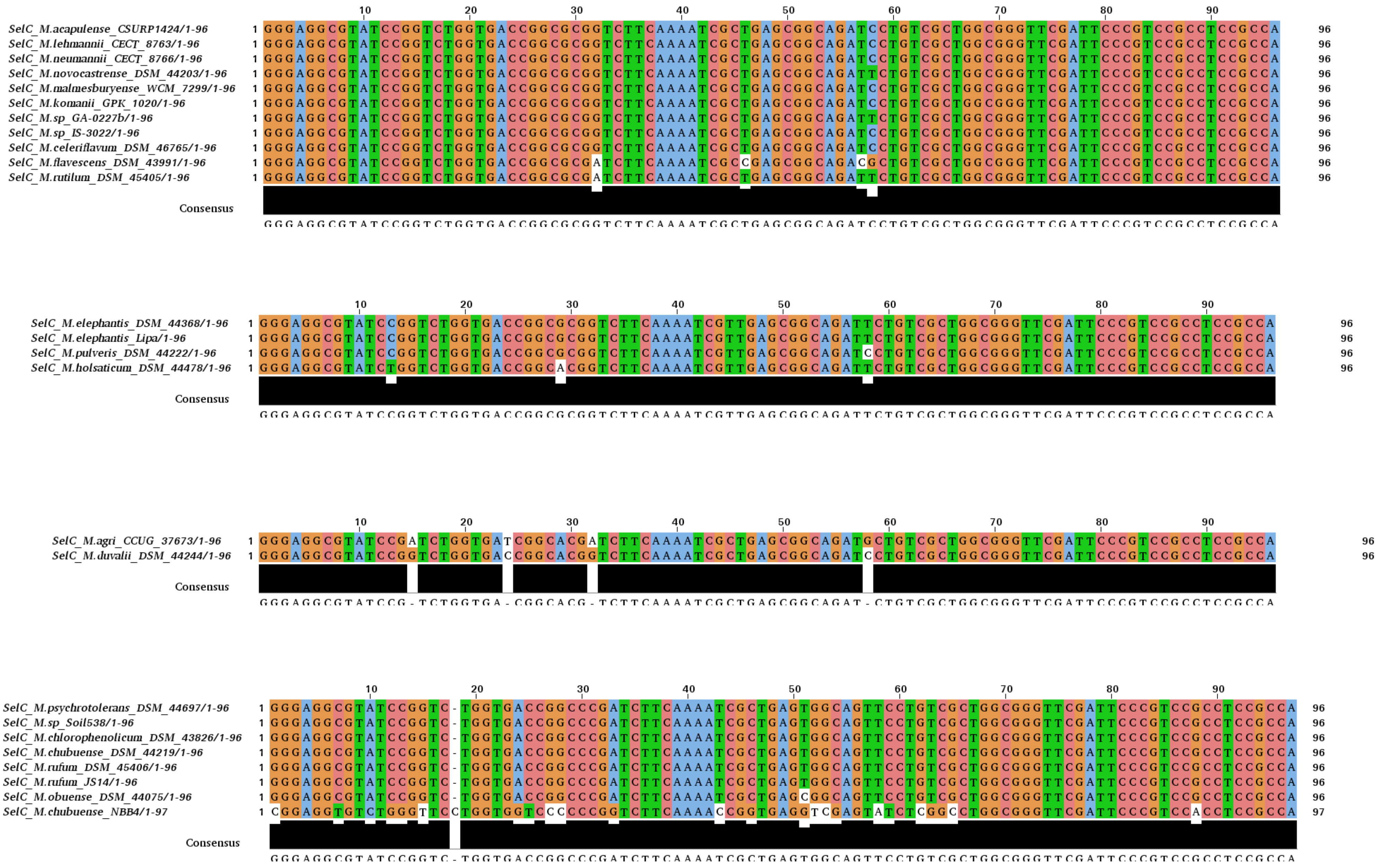

SelC_M.acapulense_CSURP1424/1-96
SelC_M.lehmannii_CECT_8763/1-96
SelC_M.neumannii_CECT_8766/1-96
SelC_M.novocastrense_DSM_44203/1-96
SelC_M.malmesburyense_WCM_7299/1-96
SelC_M.komanii_GPK_1020/1-96
SelC_M.sp_GA-0227b/1-96
SelC_M.sp_IS-3022/1-96
SelC_M.celeriflavum_DSM_46765/1-96
SelC_M.flavescens_DSM_43991/1-96
SelC_M.rutilum_DSM_45405/1-96
Consensus
GGGAGGCGTATCCGGTCTGGTGACCGGCGCGGTCTTCAAAATCGCTGAGCGGCAGATCCTGTCGCTGGCGGGTTCGATTCCCGTCCGCCTCCGCCA
SelC_M.elephantis_DSM_44368/1-96
SelC_M.elephantis_Lipa/1-96
SelC_M.pulveris_DSM_44222/1-96
SelC_M.holsaticum_DSM_44478/1-96
Consensus
GGGAGGCGTATCCGGTCTGGTGACCGGCGCGGTCTTCAAAATCGTTGAGCGGCAGATTCTGTCGCTGGCGGGTTCGATTCCCGTCCGCCTCCGCCA
SelC_M.agri_CCUG_37673/1-96
SelC_M.duvalii_DSM_44244/1-96
Consensus
GGGAGGCGTATCCG-TCTGGTGA-CGGCACG-TCTTCAAAATCGCTGAGCGGCAGAT-CTGTCGCTGGCGGGTTCGATTCCCGTCCGCCTCCGCCA
SelC_M.psychrotolerans_DSM_44697/1-96
SelC_M.sp_Soil538/1-96
SelC_M.chlorophenolicum_DSM_43826/1-96
SelC_M.chubuense_DSM_44219/1-96
SelC_M.rufum_DSM_45406/1-96
SelC_M.rufum_JS14/1-96
SelC_M.obuense_DSM_44075/1-96
SelC_M.chubuense_NBB4/1-97
Consensus
GGGAGGCGTATCCGGTC-TGGTGACCGGCCCGATCTTCAAAATCGCTGAGTGGCAGTTCCTGTCGCTGGCGGGTTCGATTCCCGTCCGCCTCCGCCA

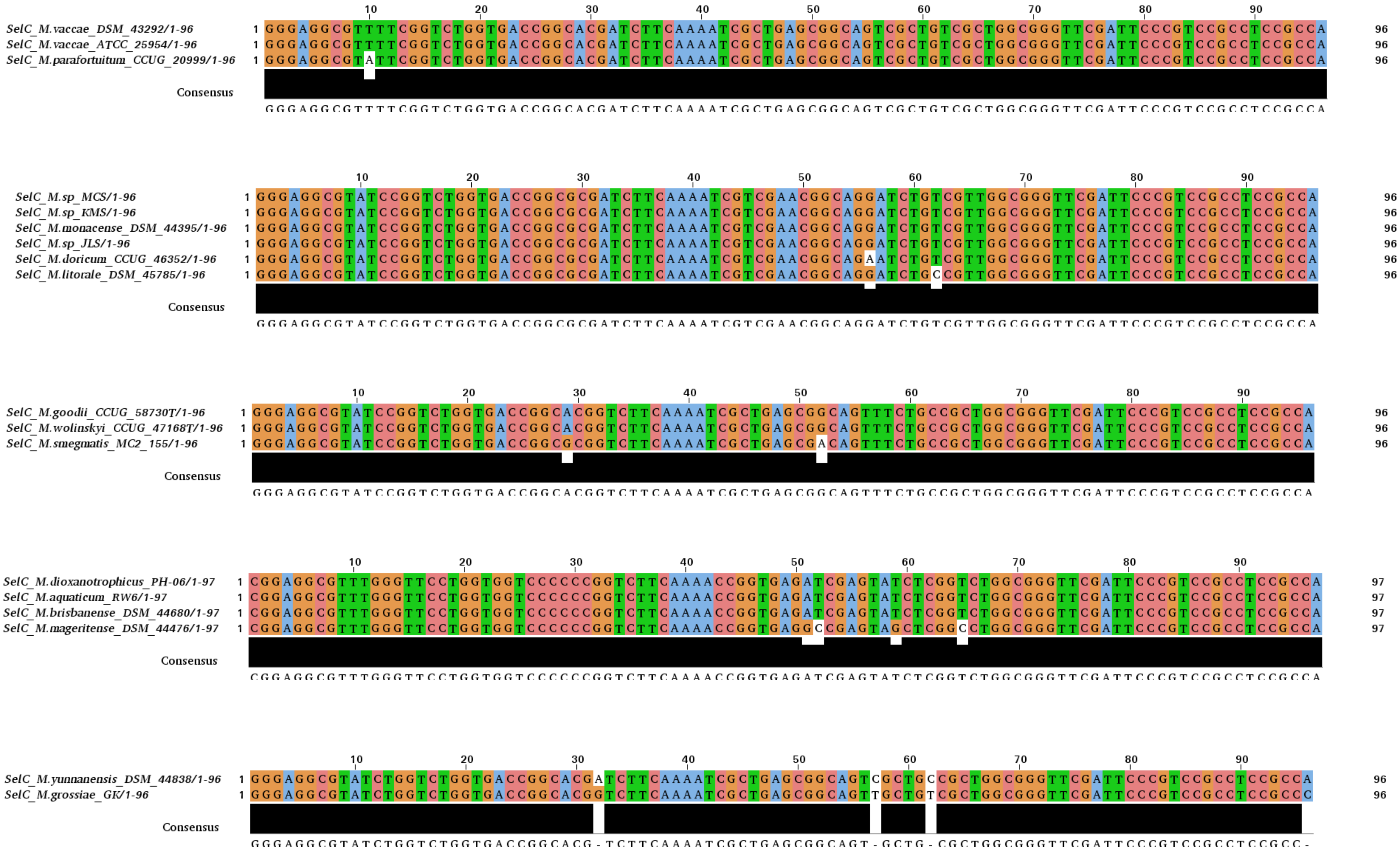
SelC_M.vaccae_DSM_43292/1-96
SelC_M.vaccae_ATCC_25954/1-96
SelC_M.parafortuitum_CCUG_20999/1-96
Consensus
GGGAGGCGTTTTCGGTCTGGTGACCGGCACGATCTTCAAAATCGCTGAGCGGCAGTCGCTGTCGCTGGCGGGTTCGATTCCCGTCCGCCTCCGCCA
SelC_M.sp_MCS/1-96
SelC_M.sp_KMS/1-96
SelC_M.monacense_DSM_44395/1-96
SelC_M.sp_JLS/1-96
SelC_M.doricum_CCUG_46352/1-96
SelC_M.litorale_DSM_45785/1-96
Consensus
GGGAGGCGTATCCGGTCTGGTGACCGGCGCGATCTTCAAAATCGTCGAACGGCAGGATCTGTCGTTGGCGGGTTCGATTCCCGTCCGCCTCCGCCA
SelC_M.goodii_CCUG_58730T/1-96
SelC_M.wolinskyi_CCUG_47168T/1-96
SelC_M.smegmatis_MC2_155/1-96
Consensus
GGGAGGCGTATCCGGTCTGGTGACCGGCACGGTCTTCAAAATCGCTGAGCGGCAGTTTCTGCCGCTGGCGGGTTCGATTCCCGTCCGCCTCCGCCA
SelC_M.dioxanotrophicus_PH-06/1-97
SelC_M.aquaticum_RW6/1-97
SelC_M.brisbanense_DSM_44680/1-97
SelC_M.mageritense_DSM_44476/1-97
Consensus
CGGAGGCGTTTGGGTTCCTGGTGGTCCCCCCGGTCTTCAAAACCGGTGAGATCGAGTATCTCGGTCTGGCGGGTTCGATTCCCGTCCGCCTCCGCCA
SelC_M.yunnanensis_DSM_44838/1-96
SelC_M.grossiae_GK/1-96
Consensus
GGGAGGCGTATCTGGTCTGGTGACCGGCACG-TCTTCAAAATCGCTGAGCGGCAGT-GCTG-CGCTGGCGGGTTCGATTCCCGTCCGCCTCCGCC-

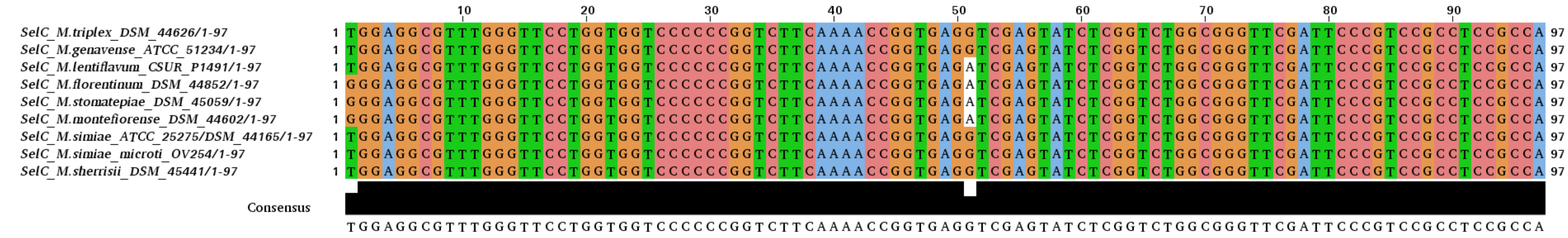
10 20 30 40 50 60 70 80 90
SelC_M.triplex_DSM_44626/1-97
SelC_M.genavense_ATCC_51234/1-97
SelC_M.lentiflavum_CSUR_P1491/1-97
SelC_M.florentinum_DSM_44852/1-97
SelC_M.stomatepiae_DSM_45059/1-97
SelC_M.montefiorense_DSM_44602/1-97
SelC_M.simiae_ATCC_25275/DSM_44165/1-97
SelC_M.simiae_microti_OV254/1-97
SelC_M.sherrisii_DSM_45441/1-97
Consensus
TGGAGGCGTTTGGGTTCCTGGTGGTCCCCCCGGTCTTCAAAACCGGTGAGGTCGAGTATCTCGGTCTGGCGGGTTCGATTCCCGTCCGCCTCCGCCA

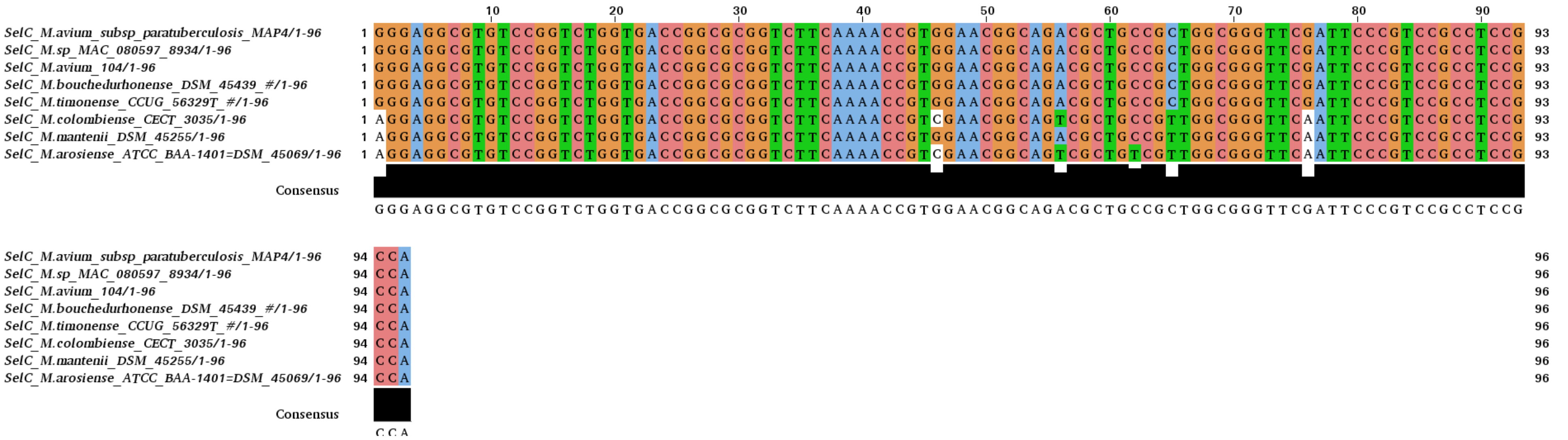
10 20 30 40 50 60 70 80 90
SelC_M.avium_subsp_paratuberculosis_MAP4/1-96
SelC_M.sp_MAC_080597_8934/1-96
SelC_M.avium_104/1-96
SelC_M.bouchedurhonense_DSM_45439_#/1-96
SelC_M.timonense_CCUG_56329T_#/1-96
SelC_M.colombiense_CECT_3035/1-96
SelC_M.mantenii_DSM_45255/1-96
SelC_M.arosiense_ATCC_BAA-1401=DSM_45069/1-96
Consensus
GGGAGGCGTGTCCGGTCTGGTGACCGGCGCGGTCTTCAAAACCGTGGAACGGCAGACGCTGCCGCTGGCGGGTTCGATTCCCGTCCGCCTCCG
CCA

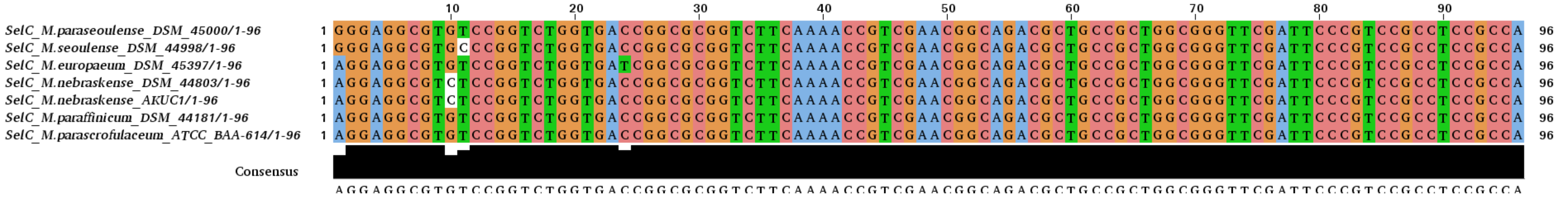
10 20 30 40 50 60 70 80 90
SelC_M.paraseoulense_DSM_45000/1-96
SelC_M.seoulense_DSM_44998/1-96
SelC_M.europaeum_DSM_45397/1-96
SelC_M.nebraskense_DSM_44803/1-96
SelC_M.nebraskense_AKUC1/1-96
SelC_M.paraffinicum_DSM_44181/1-96
SelC_M.parascrofulaceum_ATCC_BAA-614/1-96
Consensus
AGGAGGCGTGTCCGGTCTGGTGACCGGCGCGGTCTTCAAAACCGTCGAACGGCAGACGCTGCCGCTGGCGGGTTCGATTCCCGTCCGCCTCCGCCA

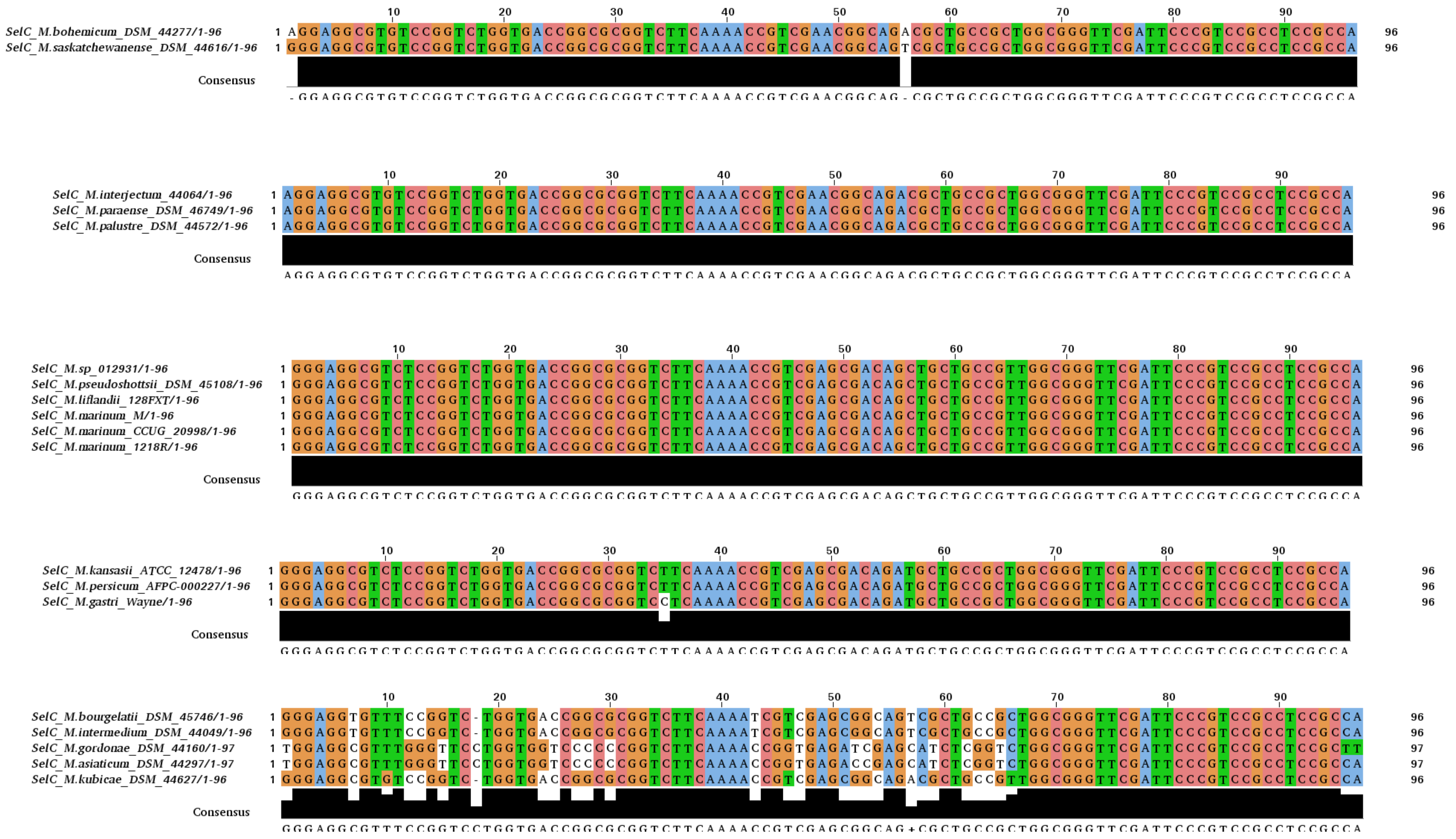
SelC_M.bohemicum_DSM_44277/1-96
SelC_M.saskatchewanense_DSM_44616/1-96
Consensus
- GGAGGCGTGTCCGGTCTGGTGACCGGCGCGGTCTTCAAAACCGTCGAACGGCAG - CGCTGCCGCTGGCGGGTTCGATTCCCGTCCGCCTCCGCCA
SelC_M.interjectum_44064/1-96
SelC_M.paraense_DSM_46749/1-96
SelC_M.palustre_DSM_44572/1-96
Consensus
AGGAGGCGTGTCCGGTCTGGTGACCGGCGCGGTCTTCAAAACCGTCGAACGGCAGACGCTGCCGCTGGCGGGTTCGATTCCCGTCCGCCTCCGCCA
SelC_M.sp_012931/1-96
SelC_M.pseudoshottsii_DSM_45108/1-96
SelC_M.liflandii_128FXT/1-96
SelC_M.marinum_M/1-96
SelC_M.marinum_CCUG_20998/1-96
SelC_M.marinum_1218R/1-96
Consensus
GGGAGGCGTCTCCGGTCTGGTGACCGGCGCGGTCTTCAAAACCGTCGAGCGACAGCTGCTGCCGTTGGCGGGTTCGATTCCCGTCCGCCTCCGCCA
SelC_M.kansasii_ATCC_12478/1-96
SelC_M.persicum_AFPC-000227/1-96
SelC_M.gastri_Wayne/1-96
Consensus
GGGAGGCGTCTCCGGTCTGGTGACCGGCGCGGTCTTCAAAACCGTCGAGCGACAGATGCTGCCGCTGGCGGGTTCGATTCCCGTCCGCCTCCGCCA
SelC_M.bourgelatii_DSM_45746/1-96
SelC_M.intermedium_DSM_44049/1-96
SelC_M.gordonae_DSM_44160/1-97
SelC_M.asiaticum_DSM_44297/1-97
SelC_M.kubicae_DSM_44627/1-96
Consensus
GGGAGGCGTTTCCGGTCCTGGTGACCGGCGCGGTCTTCAAAACCGTCGAGCGGCAG+CGCTGCCGCTGGCGGGTTCGATTCCCGTCCGCCTCCGCCA

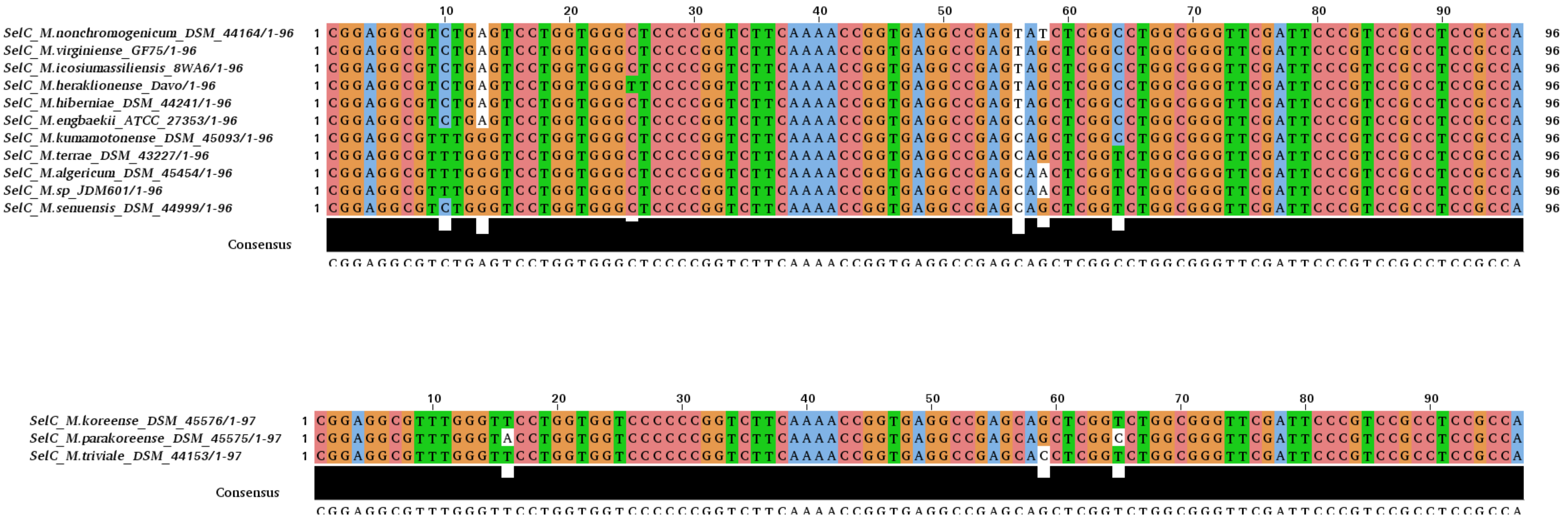
SelC_M.nonchromogenicum_DSM_44164/1-96
SelC_M.virginiense_GF75/1-96
SelC_M.icosiumassiliensis_8WA6/1-96
SelC_M.heraklionense_Davo/1-96
SelC_M.hiberniae_DSM_44241/1-96
SelC_M.engbaekii_ATCC_27353/1-96
SelC_M.kumamotonense_DSM_45093/1-96
SelC_M.terrae_DSM_43227/1-96
SelC_M.algericum_DSM_45454/1-96
SelC_M.sp_JDM601/1-96
SelC_M.senuensis_DSM_44999/1-96
Consensus
CGGAGGCGTCTGAGTCCTGGTGGGCTCCCCGGTCTTCAAAACCGGTGAGGCCGAGCAGCTCGGCCTGGCGGGTTCGATTCCCGTCCGCCTCCGCCA
SelC_M.koreense_DSM_45576/1-97
SelC_M.parakoreense_DSM_45575/1-97
SelC_M.triviale_DSM_44153/1-97
Consensus
CGGAGGCGTTTGGGTTCCTGGTGGTCCCCCCGGTCTTCAAAACCGGTGAGGCCGAGCAGCTCGGTCTGGCGGGTTCGATTCCCGTCCGCCTCCGCCA

**Figure S2**

(a) Gene synteny plot representing the chromosomal loci encompassing the SeC machinery and *fdhA* genes in different mycobacteria - *selD* (colored in red), *selC* (colored in gold), hypothetical gene(s) (colored in grey), *selA* (colored in green), *selB* (colored in turquoise), the SECIS element (colored in pink) and *fdhA* (colored in blue). The different mycobacteria are grouped according to clades as indicated to the left and a size marker is shown at the bottom to right. The gene syntenies were generated using the genoPlotR v0.8.9 R-package [46].

(b) The number of nucleotides, lower case and colored in red, between the 3' CCA end of $tRNA^{SeC}$ and the SelA translational start codon in mycobacteria in which the $tRNA^{SeC}$ gene is located immediately upstream of *selA* (vary between 11 and 75 nts). Grey shaded nucleotides correspond to putative ribosomal binding sites (Shine-Dalgarno sequence, vary between 3 and 6 nts). The different mycobacteria are grouped in clades, see main Figure 1.

Figure S2a

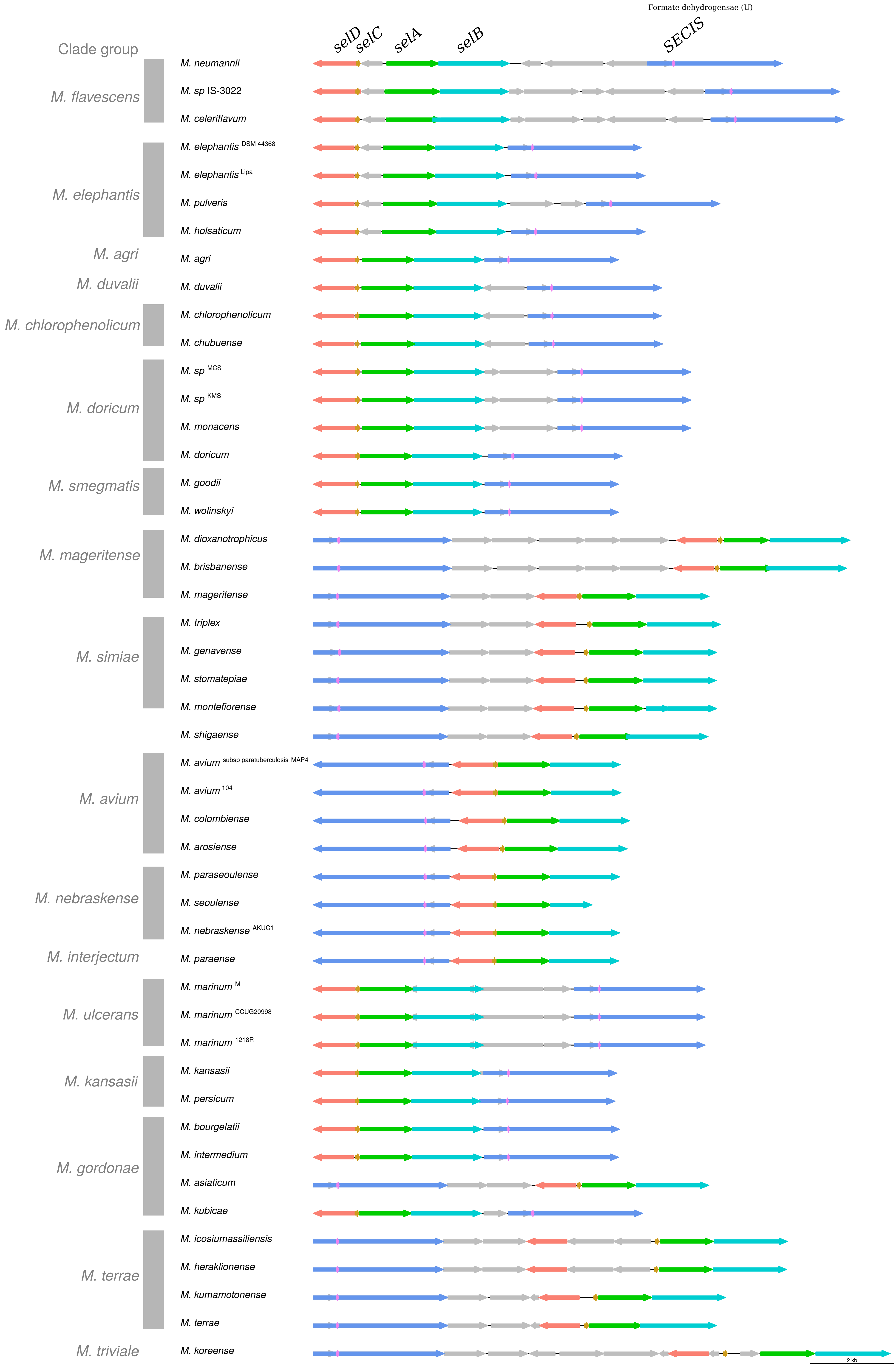
Formate dehydrogensae (U)
selD
selC
selA
selB
SECIS
Clade group
M. flavescens
M. elephantis
M. agri
M. duvalii
M. chlorophenolicum
M. doricum
M. smegmatis
M. mageritense
M. simiae
M. avium
M. nebraskense
M. interjectum
M. ulcerans
M. kansasii
M. gordonae
M. terrae
M. triviale
M. neumannii
M. sp IS-3022
M. celeriflavum
M. elephantis DSM 44368
M. elephantis Lipa
M. pulveris
M. holsaticum
M. agri
M. duvalii
M. chlorophenolicum
M. chubuense
M. sp MCS
M. sp KMS
M. monacens
M. doricum
M. goodii
M. wolinskyi
M. dioxanotrophicus
M. brisbanense
M. mageritense
M. triplex
M. genavense
M. stomatepiae
M. montefiorense
M. shigaense
M. avium subsp paratuberculosis MAP4
M. avium 104
M. colombiense
M. arosiense
M. paraseoulense
M. seoulense
M. nebraskense AKUC1
M. paraense
M. marinum M
M. marinum CCUG20998
M. marinum 1218R
M. kansasii
M. persicum
M. bourgelatii
M. intermedium
M. asiaticum
M. kubicae
M. icosiumassiliensis
M. heraklionense
M. kumamotonense
M. terrae
M. koreense
2 kb

### Figure S2b

***M. agri* clade**

>Magri SelC
…GTTCGATTCCCGTCCGCCTCCGCCAcccttcccgcgatttcggcgcgtcagggtgcgctcagcgcccgt
ttgcgcgccgaaatcgccgccgggaggcgcgGTG 75

***M. duvalii* clade**

>Mduvalii SelC
…GTTCGATTCCCGTCCGCCTCCGCCAcccctccccccgcgatttcggcgtggttatcgacgctgagcgca
cacaaacacgccgaaatcgctATG 65

***M. chlorophenolicum* clade**

>MspSoil538 SelC
…GTTCGATTCCCGTCCGCCTCCGCCAccgtccgtacgggctgcgccATG 20

>Mchloropenolicum DSM 43826 SelC
…GTTCGATTCCCGTCCGCCTCCGCCAccgtcacccctGTG 11

>Mchubuense DSM 44219 SelC
…GTTCGATTCCCGTCCGCCTCCGCCAccgttcccccctgcgaatcgggcgcgcaaaacgacagtcagc
gtcggttggcgcgccggaatcgccATG 65

>Mchubuense NBB4 SelC
…GTTCGATTCCCGTCCACCTCCGCCAtgtgtgcggcaccgaggagtagcaGTG 24

***M. austroafricanum* clade**

>Mparafortuitum CCUG 20999 SelC
…GTTCGATTCCCGTCCGCCTCCGCCAtattccacgacgaattcgccc 21

***M. doricum* clade**

>Mlitorale DSM 45785 SelC
…GTTCGATTCCCGTCCGCCTCCGCCAtactccctgcgatttcggtgtagttggttgcgttgggcgcaacca
actacaccgaaatccccgaggggccgcgccGTG 75

>Mdoricum CCUG 46352 SelC
…GTTCGATTCCCGTCCGCCTCCGCCAtaccgtccgcaaggggggcgggGTG 21

> Mmonacense DSM 44395 SelC
…GTTCGATTCCCGTCCGCCTCCGCCAtcgttcttgtgatttcggcgtgctgggtcgcgcagggcgcgaccg
cgcacgccgaaatcacgtgggtgggcgATG 72

***M. smegmatis* clade**

>Mgoodii CCUG 58730T SelC
…GTTCGATTCCCGTCCGCCTCCGCCAtattgctcctcgcgtccgcctccgccaaattgctcacccccacc
aaccacATG 51

>Mwolinskyi CCUG 47168T SelC
…GTTCGATTCCCGTCCGCCTCCGCCAtattgctcctcgcgtccgcctccgccaaattgctcacccccacc
aaccacATG 51

>Msmegmatis MC2 155 SelC
…GTTCGATTCCCGTCCGCCTCCGCCAtcgaatgccaggaccgctcctccATG 23

***M. mageritense* clade**

>Mmageritense DSM 44476 SelC
…GTTCGATTCCCGTCCGCCTCCGCCAgtcgttgctgtcggcgtcagaggagggcacGTG 30

>Mbrisbanense DSM 44680 SelC
…GTTCGATTCCCGTCCGCCTCCGCCAgtttccgttgtcgacgctcgaggagggcagGTG 30

>Maquaticum RW6 SelC
…GTTCGATTCCCGTCCGCCTCCGCCAgtttccgctgtcgacgctcgaggagggcagGTG 30

>Mdioxanotrophicus PH-06 SelC
…GTTCGATTCCCGTCCGCCTCCGCCAgtttctgctgtcgacgctcgaggagggcagGTG 30

***M. sediminis* clade**

>Myunnanensis DSM 4444838 SelC
…GTTCGATTCCCGTCCGCCTCCGCCAtgaaccacgccatggacttcgtcatgactgacccccgccgccg
ggttccccgcaccgacgcgctgcttgccgacccgcggctcgtcgacgcgcagcgactgctcggccgcacgctggtcaag
gcggtgatcgccgacgcgcaacagctcgcccgcgccggtgacatcgcgcccgacgacgtcgccgaccacgccgtcgcc
gcgctgcccgtcagcgccgccagtctcaagccggtgatcaacgcaacgggtgtcatcgtgcacaccaacctcgggcgcg
cccccctgtcccgggccgcggtcgacgcggtcgtcaccgccagcggcgccaccgacgtcgagttcgacctggccaccgg
tcgtcgcgcccgccgcggtcgcggtgcactcgcggcgcttggcgacgccgtccccacggcgcagggcgtgcacgtcgtc
aacaacaacgccgcggcgctcctgctcgccgcgatgacgctggcgcccggccgggagatcgtggtcagccgcggtgag
ctgatcgagatcggcgacgggttccggctgcctgagctgatgcagtcgacgggctcccggttccgggaggtgggcaccac
caaccgcacgcacctgcgcgactacgccgacgcgctcggtcccgacaccggcttcatcctgaaggtgcatccgtcgaact
acgtggtcagcgggttcacggccggggtgtcggtcgcggagctcgtgacgctggacgcgaccgtcgtcgtcgacgtcggct
cgggcctcttgacgccgcacccgctgctgccggaggaacccgacgcgacgacggccctgcgtgacggcgccgacctgg
tcaccgcgagcggcgacaagctgctcggcgggccgcaggccggtctgctgttcggctccgccgacctcgtcgagcggttg
cggcgccatcctgccgcgcgggcgttgcgcgtcgacaagctgacgttggccgcgctggaggcgacactggtcggaccgc
cgactccggtggcctcggcactggacgcggacgtcgcgtcactgaaggctcgcgcggtcgagttggcggggcggttgccc
ggtgccgaggccgtcgactgcgtcgccgcagtcggtggcggcggcgcaccggggatcgaattgccaagcgcggcaatc
agtttgtcggcgtcctacgcggtcgccctgcgcgccgccaggccggcggtggtcggccgcgtgcaggacggtcgatgcctc
ctcgacctgcggaccgtcgcgcccgaggacgacgaggcgctggtggcagcggtcctggcGTGta 1291

***M. simiae* clade**

>Msherrisii DSM 45441SelC
…GTTCGATTCCCGTCCGCCTCCGCCAcacgcagaccgaggaggtcgggaggtaaGTG 28

>Msimiae OV254 SelC
…GTTCGATTCCCGTCCGCCTCCGCCAgacgcagacccgaggaggtcgaggaggtaaGTG 30

>Mmontifiorense DSM 44602 SelC
…GTTCGATTCCCGTCCGCCTCCGCCAgtcgcggaccaaggaggtcgaggaggcaaGTG 29

>Mstomatepiae DSM 45059 SelC
…GTTCGATTCCCGTCCGCCTCCGCCAgacgcggacggaggaggcacGTG 20

>Mflorentium DSM 44852 SelC
…GTTCGATTCCCGTCCGCCTCCGCCAgacgcggatcgaggaggcacGTG 20

>Mlentiflavum CSUR P1491 SelC
…GTTCGATTCCCGTCCGCCTCCGCCAgccctggacgcgagcagaccgaggaggcacGTG 30

>Mgenavense ATCC 51234 SelC
…GTTCGATTCCCGTCCGCCTCCGCCAgtcgcggaccgaggaggtcgaggaaggcaaGTG 30

>Mtriplex DSM 44626 SelC
…GTTCGATTCCCGTCCGCCTCCGCCAgccgcgcacccgaggaggtcgaggaggcaaGTG 30

**Single clade**

>Mshigaense UN-152 SelC
…GTTCGATTCCCGTCCGCCTCCGCCAgccgtgccgaggaggtcgaggacgtaaGTG 27

***M. nebraskense* clade**

>Meuropaeum DSM 45397 SelC
…GTTCGATTCCCGTCCGCCTCCGCCAtacccttccgagccccacGTG 18

>Mseoulense DSM 44998 SelC
…GTTCGATTCCCGTCCGCCTCCGCCAtacttcctcgggcccgacATG 18

>Mparaseoulense DSM 45000 SelC
…GTTCGATTCCCGTCCGCCTCCGCCAtactccgtcgagcccggcATG 18

>Mparafinicum DSM 44181 SelC
…GTTCGATTCCCGTCCGCCTCCGCCAtacttcgtcgagcctgacATG 18

>Mnebraskense AKUC1 SelC
…GTTCGATTCCCGTCCGCCTCCGCCAtacttcttcgagcctgcATG 17

>Mnebraskense DSM 44803 SelC
…GTTCGATTCCCGTCCGCCTCCGCCAtacttcttcgagcctgcATG 17

**Single clade**

>Mparmense DSM 44553 SelC
…GTTCGATTCCCGTCCGCCTCCGCCAtgctccctggggtgcggcATG 18

***M. bohemicum* clade**

>Mbohemicum DSM 44277 SelC
…GTTCGATTCCCGTCCGCCTCCGCCAtacttcctccggcccgacATG 18

***M. interjectum* clade**

>Mparaense DSM 46747 SelC
…GTTCGATTCCCGTCCGCCTCCGCCAtacttcgtcgagcccgagATG 18

***M. ulcerans* clade**

>Msp012931 SelC
…GTTCGATTCCCGTCCGCCTCCGCCAccgcgccaatcgccatgggtgacggcagcgaccggcgccgc
gcaccgacGTG 49

>Mpseudoshottsii DSM 45108 SelC
…GTTCGATTCCCGTCCGCCTCCGCCAccgcgccaatcgccatgggtgacggcagcgaccggcgccgc
gcaccgacGTG 49

>Mmarinum CCUG 20998 SelC
…GTTCGATTCCCGTCCGCCTCCGCCAccgcgccgatcgccATG 14

>Mliflandi 128FXT SelC
…GTTCGATTCCCGTCCGCCTCCGCCAccgcgccgatcgccATG 14

***M. kansasii* clade**

>Mpersicum AFPC-000227 SelC
…GTTCGATTCCCGTCCGCCTCCGCCAcctccctgggtccggacATG 17

>Mkansasii ATCC 12478 SelC
…GTTCGATTCCCGTCCGCCTCCGCCAcctccccgggtccagacATG 17

>Mgastri Wayne SelC
…GTTCGATTCCCGTCCGCCTCCGCCAcctccctgggtccggacATG 17

***M. gordonae* clade**

>Mintermedium DSM 44049 SelC
…GTTCGATTCCCGTCCGCCTCCGCCAccctgttacgtcATG 12

>Mkubicae DSM 44627 SelC
…GTTCGATTCCCGTCCGCCTCCGCCAccctggtacctgagATG 14

>Masiaticum DSM 44297 SelC
…GTTCGATTCCCGTCCGCCTCCGCCAgttgcctacctagggaagcggtgaagatcaGTG 30

>Mbourgelatii DSM 45746 SelC
…GTTCGATTCCCGTCCGCCTCCGCCAccctggtaccccATG 12

>Mgordonae DSM 44160 SelC
…GTTCGATTCCCGTCCGCCTCCGCttttgcttcaaaagtgaaagcccaccgagagaggtcacGTG 38

***M. terrae* clade**
>Mterrae DSM 43227 SelC
…GTTCGATTCCCGTCCGCCTCCGCCAtaggaggcgctGTG 11

>MspJDM601 SelC
…GTTCGATTCCCGTCCGCCTCCGCCAtcggacaggagggggctGTG 16

>Msenuensis DSM 44999 SelC
…GTTCGATTCCCGTCCGCCTCCGCCAccggacaggagggggctGTG 16

>Mkumamotonense DSM 45093 SelC
…GTTCGATTCCCGTCCGCCTCCGCCAtccgacaggaggcgcaGTG 16

>Malgericum DSM 45454 SelC
…GTTCGATTCCCGTCCGCCTCCGCCAtcggacaggagggggctGTG 16

>Mengbaekii ATCC 27353 SelC
…GTTCGATTCCCGTCCGCCTCCGCCAgctgcacatccaggaggagctGTG 21

>Mheraklionense Davo SelC
…GTTCGATTCCCGTCCGCCTCCGCCAgctgcgtatcgccaggaggagccGTG 23

>Mhiberniae DSM 44241 SelC
…GTTCGATTCCCGTCCGCCTCCGCCAgctgcacatccaggaggagctGTG 21

>Micosiumassiliense 8WA6 SelC
…GTTCGATTCCCGTCCGCCTCCGCCAgctgagtatctccaggaggagctGTG 23

>Mnonchromogenicum DSM 44164 SelC
…GTTCGATTCCCGTCCGCCTCCGCCAgctgcgtatcggaggagaagctGTG 22

***M. triviale* clade**

>Mparakoreense DSM 45575 SelC
…GTTCGATTCCCGTCCGCCTCCGCCAaccgcgccgaggagggttGTG 18

>Mkoreense DSM 45576 SelC
…GTTCGATTCCCGTCCGCCTCCGCCAccccgctcgaggagggttGTG 18

**Figure S3**

Alignment of the SECIS element sequences and grouped according to clades beginning with the *M. flavescens* clade and ending with the *M. triviale* clade, see main Figure 1.

Figure S3

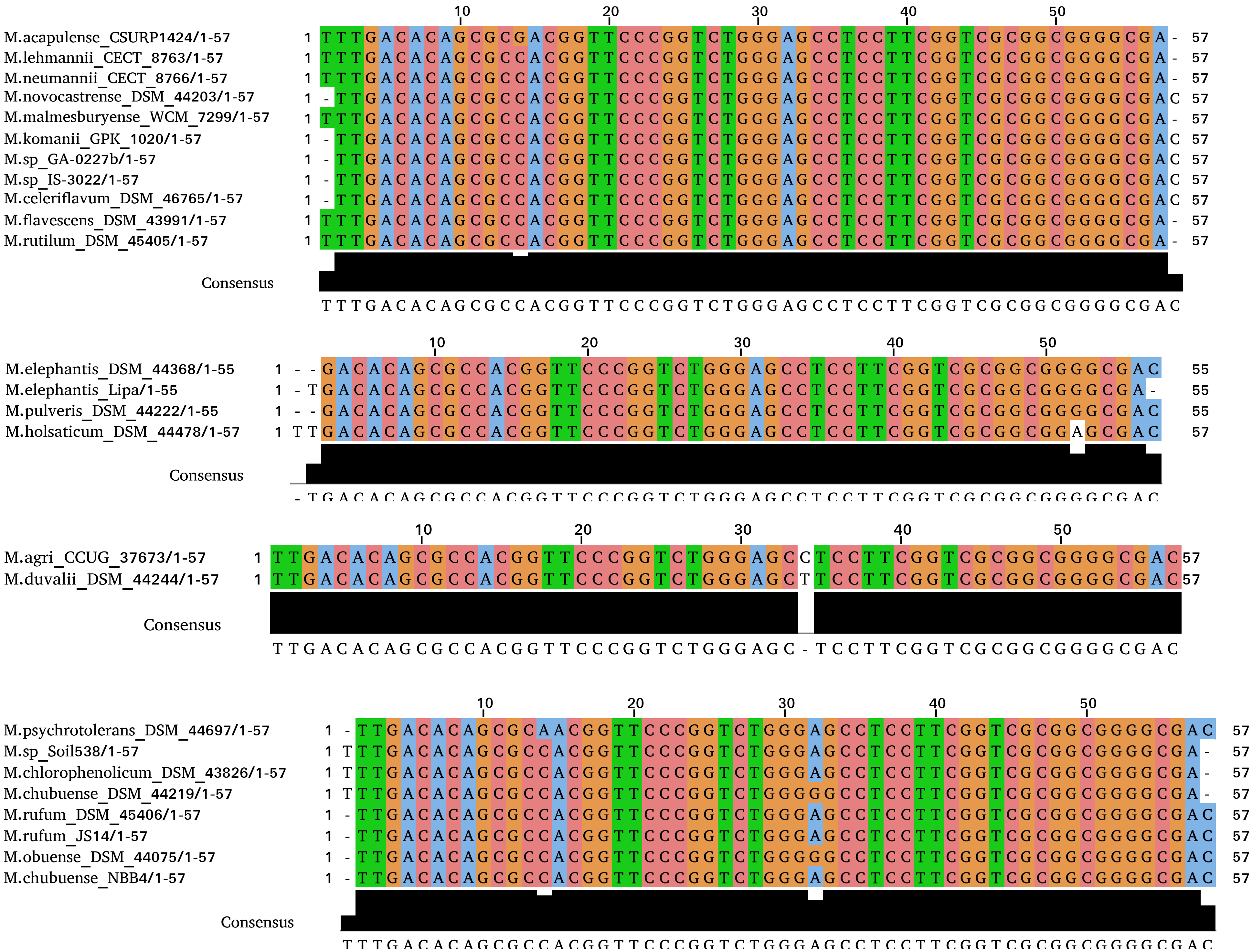
10 20 30 40 50
M.acapulense_CSURP1424/1-57 1 TTTGACACAGCGCGACGGTTCCCGGTCTGGGAGCCTCCTTCGGTCGCGGCGGGGCGA- 57
M.lehmannii_CECT_8763/1-57 1 TTTGACACAGCGCCACGGTTCCCGGTCTGGGAGCCTCCTTCGGTCGCGGCGGGGCGA- 57
M.neumannii_CECT_8766/1-57 1 TTTGACACAGCGCCACGGTTCCCGGTCTGGGAGCCTCCTTCGGTCGCGGCGGGGCGA- 57
M.novocastrense_DSM_44203/1-57 1 -TTGACACAGCGCCACGGTTCCCGGTCTGGGAGCCTCCTTCGGTCGCGGCGGGGCGAC 57
M.malmesburyense_WCM_7299/1-57 1 TTTGACACAGCGCCACGGTTCCCGGTCTGGGAGCCTCCTTCGGTCGCGGCGGGGCGA- 57
M.komanii_GPK_1020/1-57 1 -TTGACACAGCGCCACGGTTCCCGGTCTGGGAGCCTCCTTCGGTCGCGGCGGGGCGAC 57
M.sp_GA-0227b/1-57 1 -TTGACACAGCGCCACGGTTCCCGGTCTGGGAGCCTCCTTCGGTCGCGGCGGGGCGAC 57
M.sp_IS-3022/1-57 1 -TTGACACAGCGCCACGGTTCCCGGTCTGGGAGCCTCCTTCGGTCGCGGCGGGGCGAC 57
M.celeriflavum_DSM_46765/1-57 1 -TTGACACAGCGCCACGGTTCCCGGTCTGGGAGCCTCCTTCGGTCGCGGCGGGGCGAC 57
M.flavescens_DSM_43991/1-57 1 TTTGACACAGCGCCACGGTTCCCGGTCTGGGAGCCTCCTTCGGTCGCGGCGGGGCGA- 57
M.rutilum_DSM_45405/1-57 1 TTTGACACAGCGCCACGGTTCCCGGTCTGGGAGCCTCCTTCGGTCGCGGCGGGGCGA- 57
Consensus
TTTGACACAGCGCCACGGTTCCCGGTCTGGGAGCCTCCTTCGGTCGCGGCGGGGCGAC
10 20 30 40 50
M.elephantis_DSM_44368/1-55 1 --GACACAGCGCCACGGTTCCCGGTCTGGGAGCCTCCTTCGGTCGCGGCGGGGCGAC 55
M.elephantis_Lipa/1-55 1 -TGACACAGCGCCACGGTTCCCGGTCTGGGAGCCTCCTTCGGTCGCGGCGGGGCGA- 55
M.pulveris_DSM_44222/1-55 1 --GACACAGCGCCACGGTTCCCGGTCTGGGAGCCTCCTTCGGTCGCGGCGGGGCGAC 55
M.holsaticum_DSM_44478/1-57 1 TTGACACAGCGCCACGGTTCCCGGTCTGGGAGCCTCCTTCGGTCGCGGCGGAGCGAC 57
Consensus
-TGACACAGCGCCACGGTTCCCGGTCTGGGAGCCTCCTTCGGTCGCGGCGGGGCGAC
10 20 30 40 50
M.agri_CCUG_37673/1-57 1 TTGACACAGCGCCACGGTTCCCGGTCTGGGAGCCTCCTTCGGTCGCGGCGGGGCGAC 57
M.duvalii_DSM_44244/1-57 1 TTGACACAGCGCCACGGTTCCCGGTCTGGGAGCTTCCTTCGGTCGCGGCGGGGCGAC 57
Consensus
TTGACACAGCGCCACGGTTCCCGGTCTGGGAGC-TCCTTCGGTCGCGGCGGGGCGAC
10 20 30 40 50
M.psychrotolerans_DSM_44697/1-57 1 -TTGACACAGCGCAACGGTTCCCGGTCTGGGAGCCTCCTTCGGTCGCGGCGGGGCGAC 57
M.sp_Soil538/1-57 1 TTTGACACAGCGCCACGGTTCCCGGTCTGGGAGCCTCCTTCGGTCGCGGCGGGGCGA- 57
M.chlorophenolicum_DSM_43826/1-57 1 TTTGACACAGCGCCACGGTTCCCGGTCTGGGAGCCTCCTTCGGTCGCGGCGGGGCGA- 57
M.chubuense_DSM_44219/1-57 1 TTTGACACAGCGCCACGGTTCCCGGTCTGGGGGCCTCCTTCGGTCGCGGCGGGGCGA- 57
M.rufum_DSM_45406/1-57 1 -TTGACACAGCGCCACGGTTCCCGGTCTGGGAGCCTCCTTCGGTCGCGGCGGGGCGAC 57
M.rufum_JS14/1-57 1 -TTGACACAGCGCCACGGTTCCCGGTCTGGGAGCCTCCTTCGGTCGCGGCGGGGCGAC 57
M.obuense_DSM_44075/1-57 1 -TTGACACAGCGCCACGGTTCCCGGTCTGGGGGCCTCCTTCGGTCGCGGCGGGGCGAC 57
M.chubuense_NBB4/1-57 1 -TTGACACAGCGCCACGGTTCCCGGTCTGGGAGCCTCCTTCGGTCGCGGCGGGGCGAC 57
Consensus
TTTGACACAGCGCCACGGTTCCCGGTCTGGGAGCCTCCTTCGGTCGCGGCGGGGCGAC

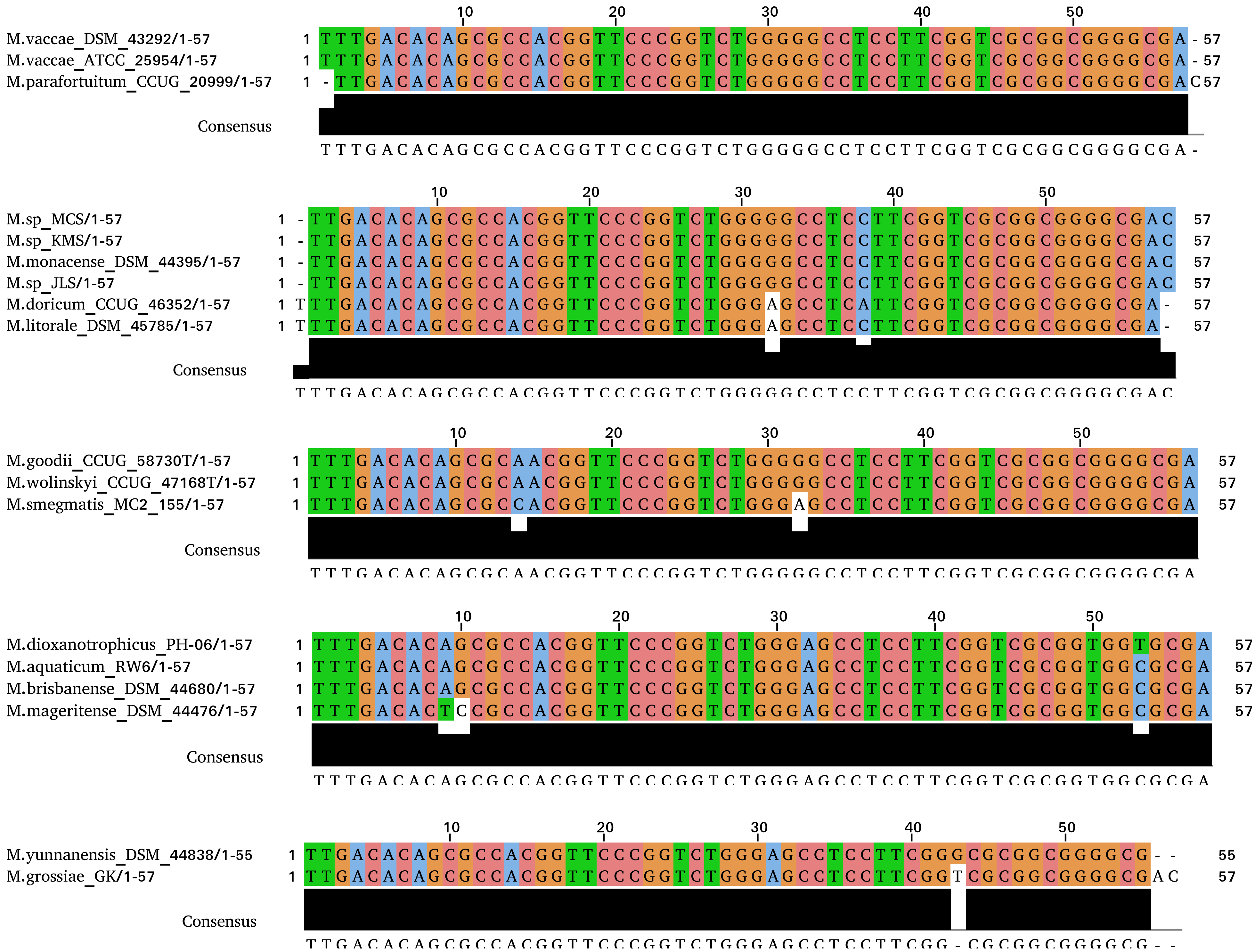
M.vaccae_DSM_43292/1-57
M.vaccae_ATCC_25954/1-57
M.parafortuitum_CCUG_20999/1-57
Consensus
T T T G A C A C A G C G C C A C G G T T C C C G G T C T G G G G G C C T C C T T C G G T C G C G G C G G G G C G A -
M.sp_MCS/1-57
M.sp_KMS/1-57
M.monacense_DSM_44395/1-57
M.sp_JLS/1-57
M.doricum_CCUG_46352/1-57
M.litorale_DSM_45785/1-57
Consensus
T T T G A C A C A G C G C C A C G G T T C C C G G T C T G G G G G C C T C C T T C G G T C G C G G C G G G G C G A C
M.goodii_CCUG_58730T/1-57
M.wolinskyi_CCUG_47168T/1-57
M.smegmatis_MC2_155/1-57
Consensus
T T T G A C A C A G C G C A A C G G T T C C C G G T C T G G G G G C C T C C T T C G G T C G C G G C G G G G C G A
M.dioxanotrophicus_PH-06/1-57
M.aquaticum_RW6/1-57
M.brisbanense_DSM_44680/1-57
M.mageritense_DSM_44476/1-57
Consensus
T T T G A C A C A G C G C C A C G G T T C C C G G T C T G G G A G C C T C C T T C G G T C G C G G T G G C G C G A
M.yunnanensis_DSM_44838/1-55
M.grossiae_GK/1-57
Consensus
T T G A C A C A G C G C C A C G G T T C C C G G T C T G G G A G C C T C C T T C G G - C G C G G C G G G G C G - -

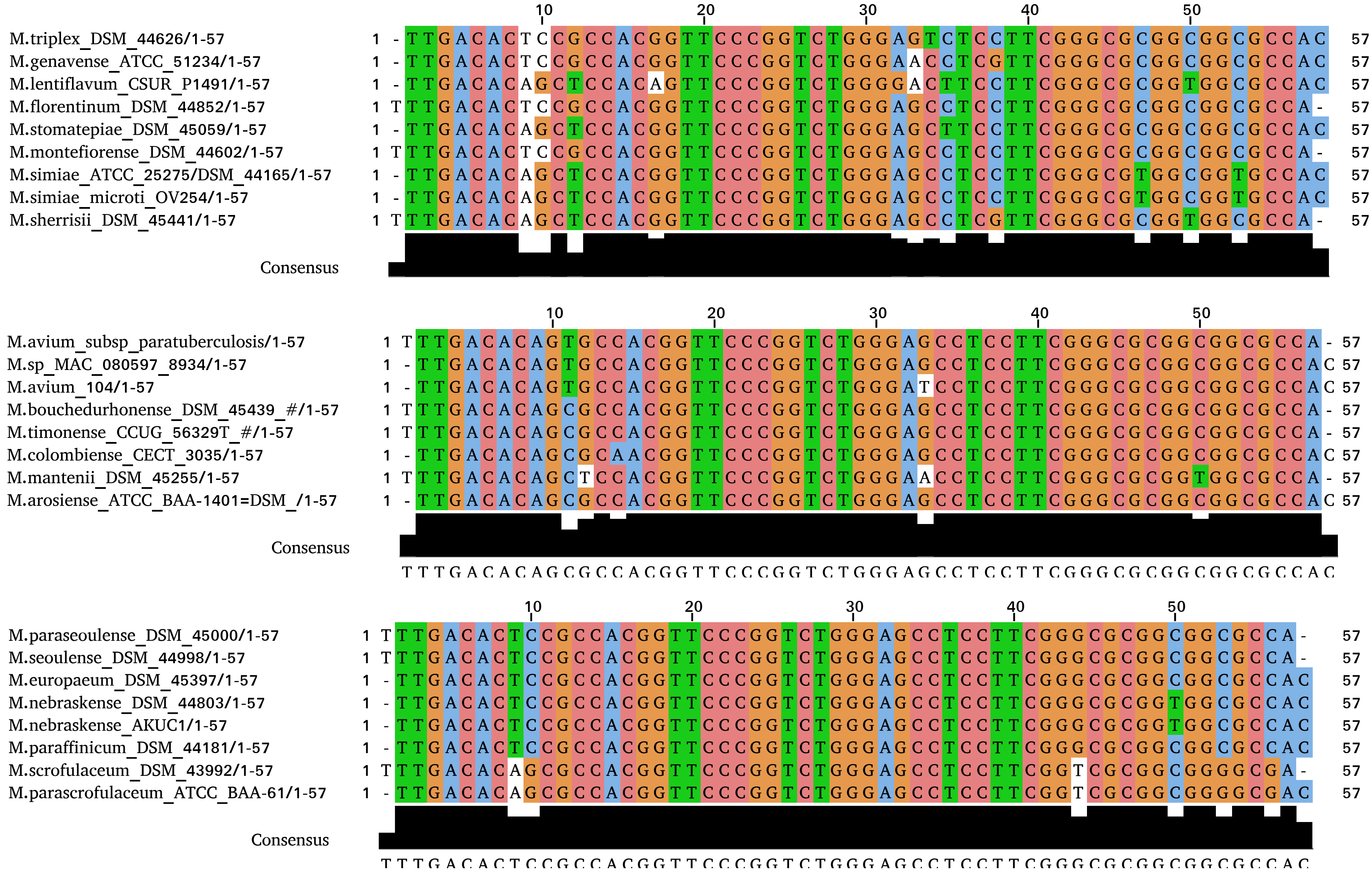
M.triplex_DSM_44626/1-57
M.genavense_ATCC_51234/1-57
M.lentiflavum_CSUR_P1491/1-57
M.florentinum_DSM_44852/1-57
M.stomatepiae_DSM_45059/1-57
M.montefiorense_DSM_44602/1-57
M.simiae_ATCC_25275/DSM_44165/1-57
M.simiae_microti_OV254/1-57
M.sherrisii_DSM_45441/1-57
Consensus
M.avium_subsp_paratuberculosis/1-57
M.sp_MAC_080597_8934/1-57
M.avium_104/1-57
M.bouchedurhonense_DSM_45439_#/1-57
M.timonense_CCUG_56329T_#/1-57
M.colombiense_CECT_3035/1-57
M.mantenii_DSM_45255/1-57
M.arosiense_ATCC_BAA-1401=DSM_/1-57
Consensus
T T T G A C A C A G C G C C A C G G T T C C C G G T C T G G G A G C C T C C T T C G G G C G C G G C G G C G C C A C
M.paraseoulense_DSM_45000/1-57
M.seoulense_DSM_44998/1-57
M.europaeum_DSM_45397/1-57
M.nebraskense_DSM_44803/1-57
M.nebraskense_AKUC1/1-57
M.paraffinicum_DSM_44181/1-57
M.scrofulaceum_DSM_43992/1-57
M.parascrofulaceum_ATCC_BAA-61/1-57
Consensus
T T T G A C A C T C C G C C A C G G T T C C C G G T C T G G G A G C C T C C T T C G G G C G C G G C G G C G C C A C

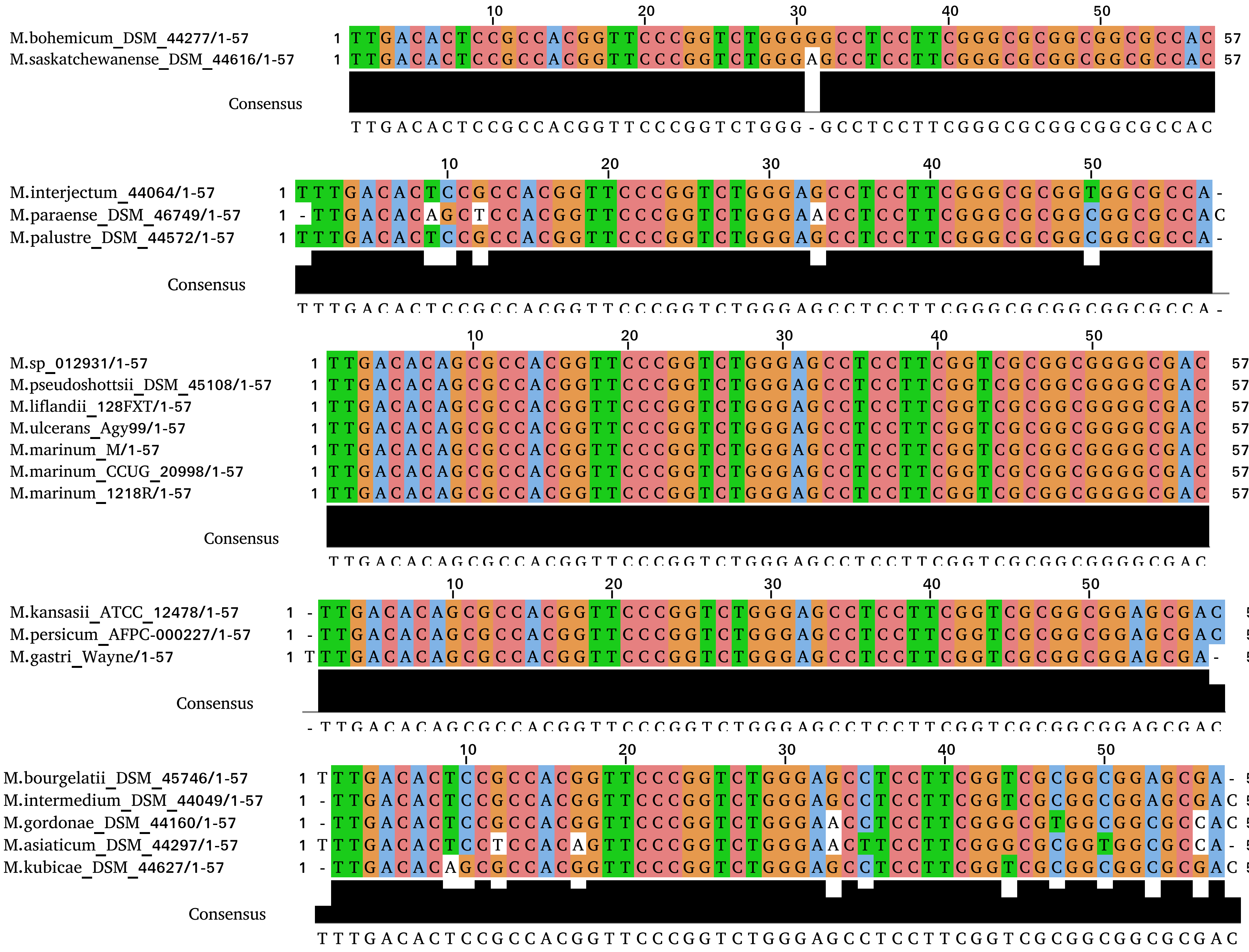
M.bohemicum_DSM_44277/1-57
M.saskatchewanense_DSM_44616/1-57
Consensus
M.interjectum_44064/1-57
M.paraense_DSM_46749/1-57
M.palustre_DSM_44572/1-57
Consensus
M.sp_012931/1-57
M.pseudoshottsii_DSM_45108/1-57
M.liflandii_128FXT/1-57
M.ulcerans_Agy99/1-57
M.marinum_M/1-57
M.marinum_CCUG_20998/1-57
M.marinum_1218R/1-57
Consensus
M.kansasii_ATCC_12478/1-57
M.persicum_AFPC-000227/1-57
M.gastri_Wayne/1-57
Consensus
M.bourgelatii_DSM_45746/1-57
M.intermedium_DSM_44049/1-57
M.gordonae_DSM_44160/1-57
M.asiaticum_DSM_44297/1-57
M.kubicae_DSM_44627/1-57
Consensus

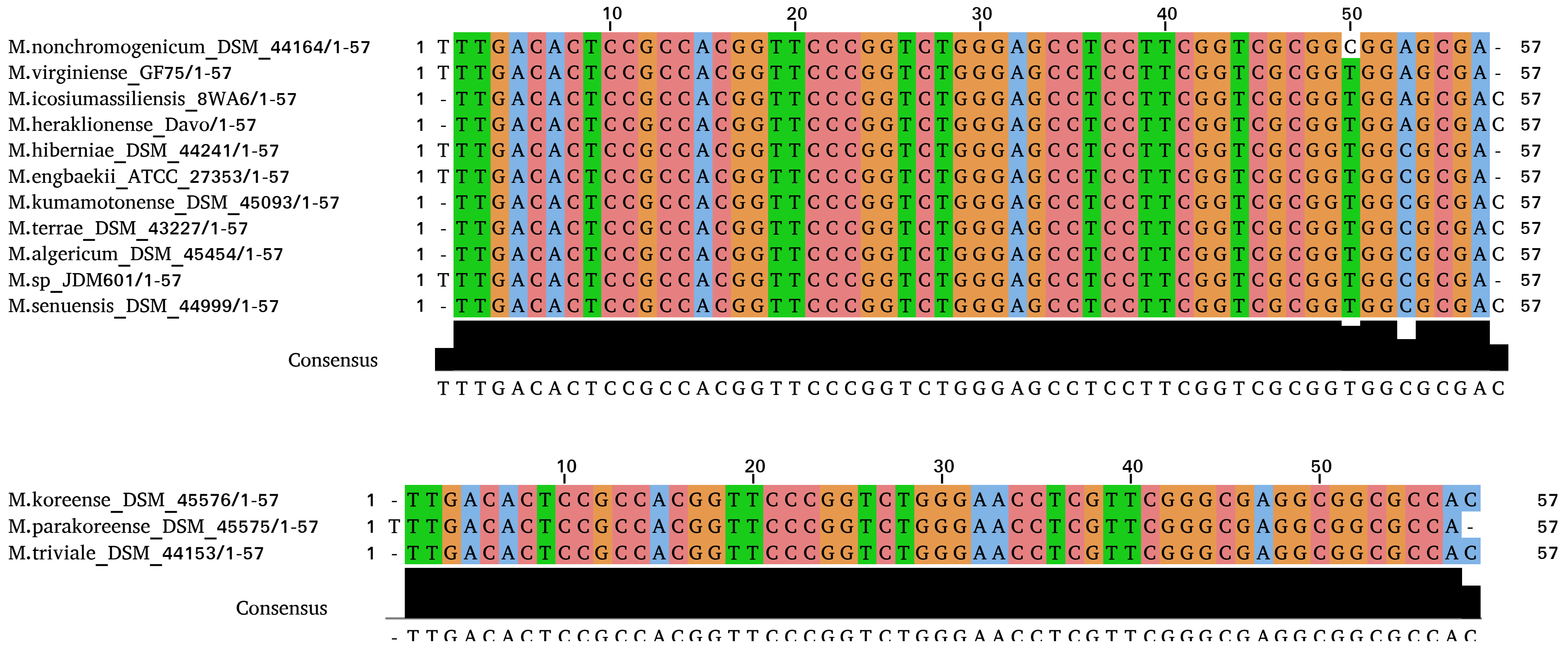
10 20 30 40 50
M.nonchromogenicum_DSM_44164/1-57 1 TTTGACACTCCGCCACGGTTCCCGGTCTGGGAGCCTCCTTCGGTCGCGGCGGAGCGA- 57
M.virginiense_GF75/1-57 1 TTTGACACTCCGCCACGGTTCCCGGTCTGGGAGCCTCCTTCGGTCGCGGTGGAGCGA- 57
M.icosiumassiliensis_8WA6/1-57 1 -TTGACACTCCGCCACGGTTCCCGGTCTGGGAGCCTCCTTCGGTCGCGGTGGAGCGAC 57
M.heraklionense_Davo/1-57 1 -TTGACACTCCGCCACGGTTCCCGGTCTGGGAGCCTCCTTCGGTCGCGGTGGAGCGAC 57
M.hiberniae_DSM_44241/1-57 1 TTTGACACTCCGCCACGGTTCCCGGTCTGGGAGCCTCCTTCGGTCGCGGTGGCGCGA- 57
M.engbaekii_ATCC_27353/1-57 1 TTTGACACTCCGCCACGGTTCCCGGTCTGGGAGCCTCCTTCGGTCGCGGTGGCGCGA- 57
M.kumamotonense_DSM_45093/1-57 1 -TTGACACTCCGCCACGGTTCCCGGTCTGGGAGCCTCCTTCGGTCGCGGTGGCGCGAC 57
M.terrae_DSM_43227/1-57 1 -TTGACACTCCGCCACGGTTCCCGGTCTGGGAGCCTCCTTCGGTCGCGGTGGCGCGAC 57
M.algericum_DSM_45454/1-57 1 -TTGACACTCCGCCACGGTTCCCGGTCTGGGAGCCTCCTTCGGTCGCGGTGGCGCGAC 57
M.sp_JDM601/1-57 1 TTTGACACTCCGCCACGGTTCCCGGTCTGGGAGCCTCCTTCGGTCGCGGTGGCGCGA- 57
M.senuensis_DSM_44999/1-57 1 -TTGACACTCCGCCACGGTTCCCGGTCTGGGAGCCTCCTTCGGTCGCGGTGGCGCGAC 57
Consensus
TTTGACACTCCGCCACGGTTCCCGGTCTGGGAGCCTCCTTCGGTCGCGGTGGCGCGAC
10 20 30 40 50
M.koreense_DSM_45576/1-57 1 -TTGACACTCCGCCACGGTTCCCGGTCTGGGAACCTCGTTCGGGCGAGGCGGCGCCAC 57
M.parakoreense_DSM_45575/1-57 1 TTTGACACTCCGCCACGGTTCCCGGTCTGGGAACCTCGTTCGGGCGAGGCGGCGCCA- 57
M.triviale_DSM_44153/1-57 1 -TTGACACTCCGCCACGGTTCCCGGTCTGGGAACCTCGTTCGGGCGAGGCGGCGCCAC 57
Consensus
-TTGACACTCCGCCACGGTTCCCGGTCTGGGAACCTCGTTCGGGCGAGGCGGCGCCAC

**Figure S4**

Phylogenetic trees for SeC-machinery and *fdhA* genes. Mycobacteria colored in green and red refer to RGM and SGM, respectively. The phylogenetic trees are based on MAFFT multiple sequence alignment computed using the FastTree (v2.1.19) [35] and 1000 cycles of bootstrapping and default settings. The FastTree infers approximately-maximum-likelihood phylogenetic trees from protein sequence alignments. For comparison, we included *Rhodococcus hoagie*, *Rhodococcus equi*, *Corynebacterium massiliense*, *Tsukamurella* sp. 1534 and *Gordonia bronchialis*, which also have the SeC-machinery and FdhA genes. The different mycobacteria were grouped into two main groups 1 and 2, and group 2 was divided into 2.1 and 2.2 as indicated to the right.

(a) SelA

(b) SelB

(c) $tRNA^{SeC}$ (the different mycobacteria were not colored according to growth)

(d) SelD

(e) FdhA

### Figure S4a

SelA

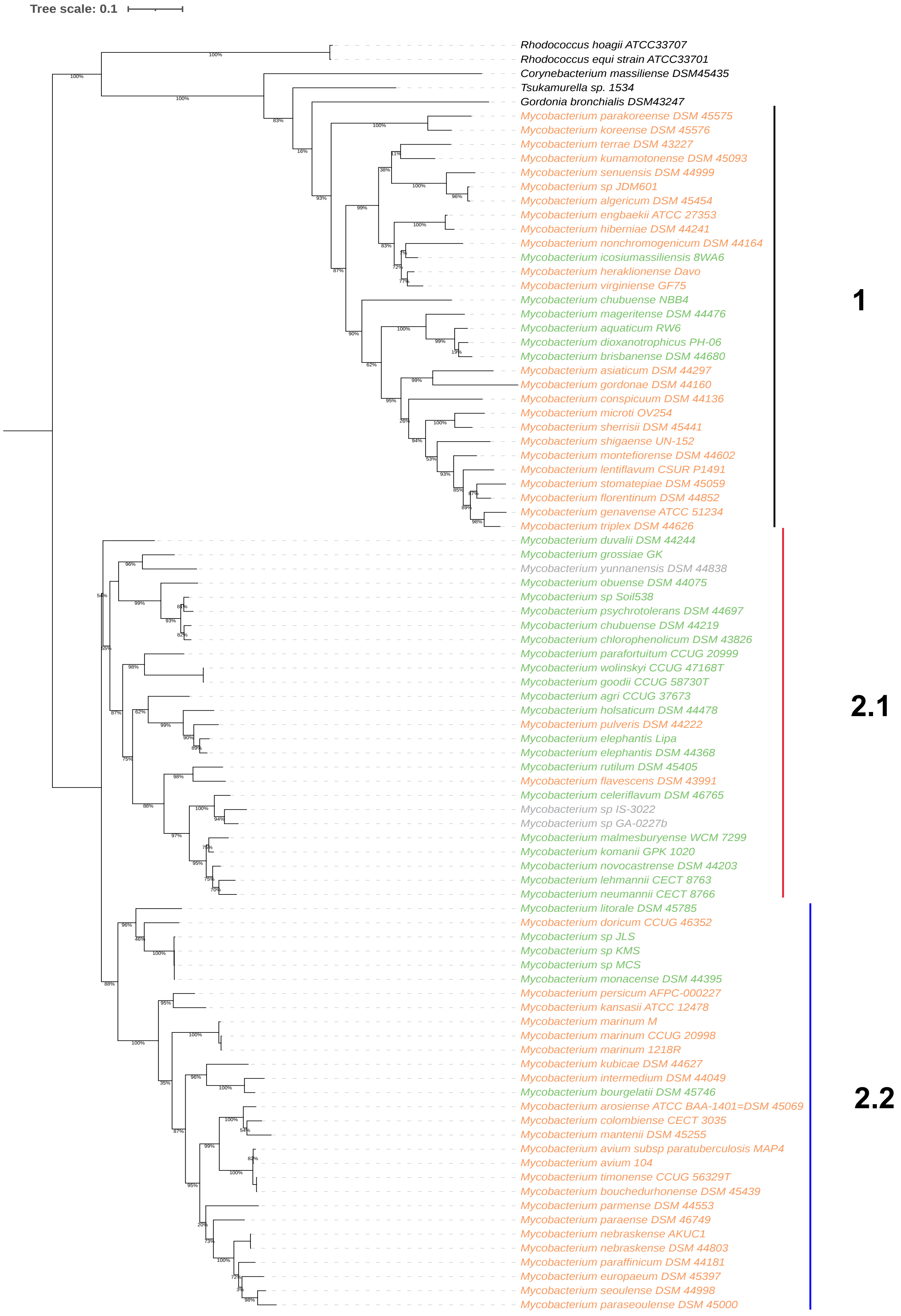

### Figure S4b

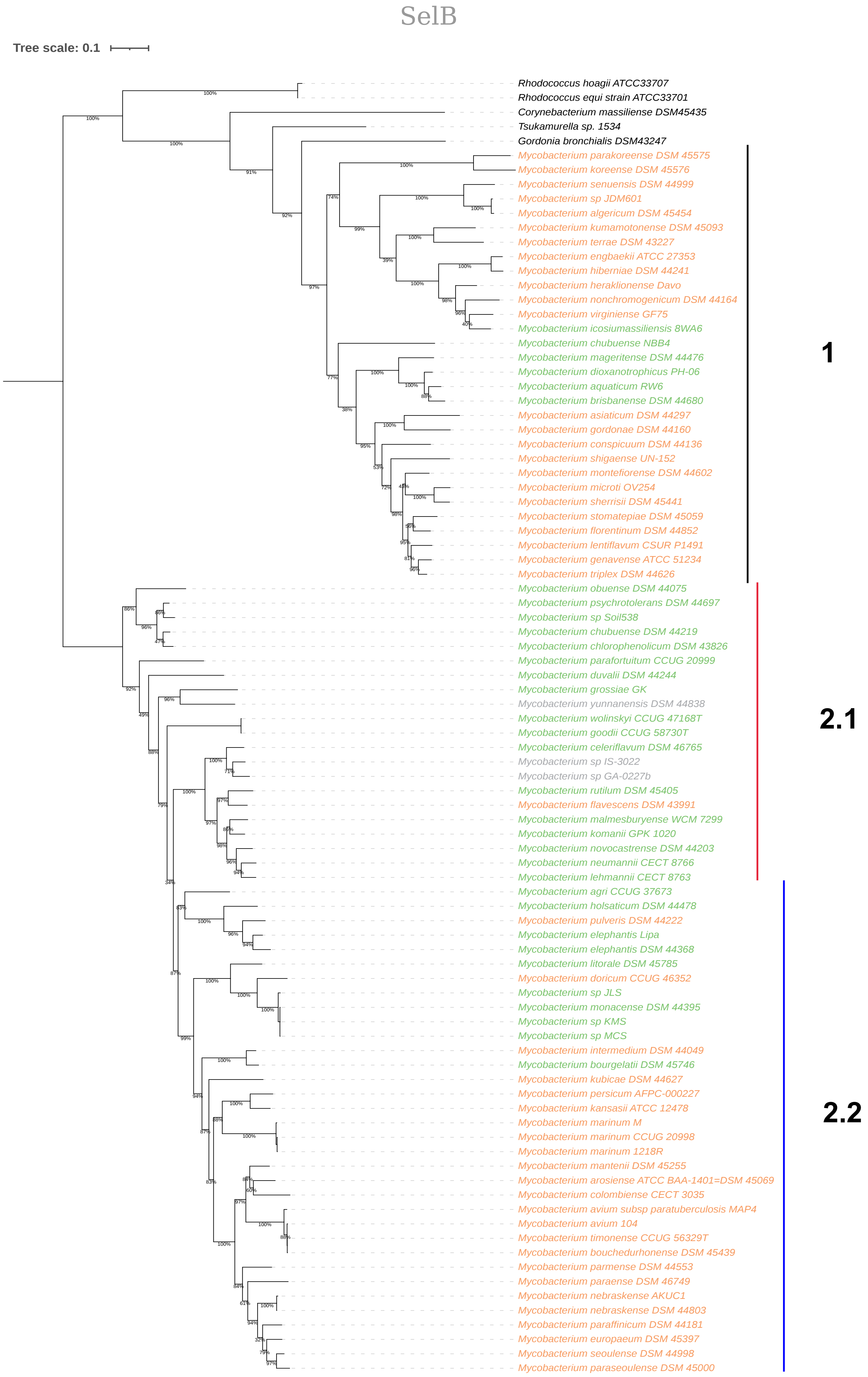
SelB
Tree scale: 0.1
Rhodococcus hoagii ATCC33707
Rhodococcus equi strain ATCC33701
Corynebacterium massiliense DSM45435
Tsukamurella sp. 1534
Gordonia bronchialis DSM43247
Mycobacterium parakoreense DSM 45575
Mycobacterium koreense DSM 45576
Mycobacterium senuensis DSM 44999
Mycobacterium sp JDM601
Mycobacterium algericum DSM 45454
Mycobacterium kumamotonense DSM 45093
Mycobacterium terrae DSM 43227
Mycobacterium engbaekii ATCC 27353
Mycobacterium hiberniae DSM 44241
Mycobacterium heraklionense Davo
Mycobacterium nonchromogenicum DSM 44164
Mycobacterium virginiense GF75
Mycobacterium icosiumassiliensis 8WA6
Mycobacterium chubuense NBB4
Mycobacterium mageritense DSM 44476
Mycobacterium dioxanotrophicus PH-06
Mycobacterium aquaticum RW6
Mycobacterium brisbanense DSM 44680
Mycobacterium asiaticum DSM 44297
Mycobacterium gordonae DSM 44160
Mycobacterium conspicuum DSM 44136
Mycobacterium shigaense UN-152
Mycobacterium montefiorense DSM 44602
Mycobacterium microti OV254
Mycobacterium sherrisii DSM 45441
Mycobacterium stomatepiae DSM 45059
Mycobacterium florentinum DSM 44852
Mycobacterium lentiflavum CSUR P1491
Mycobacterium genavense ATCC 51234
Mycobacterium triplex DSM 44626
Mycobacterium obuense DSM 44075
Mycobacterium psychrotolerans DSM 44697
Mycobacterium sp Soil538
Mycobacterium chubuense DSM 44219
Mycobacterium chlorophenolicum DSM 43826
Mycobacterium parafortuitum CCUG 20999
Mycobacterium duvalii DSM 44244
Mycobacterium grossiae GK
Mycobacterium yunnanensis DSM 44838
Mycobacterium wolinskyi CCUG 47168T
Mycobacterium goodii CCUG 58730T
Mycobacterium celeriflavum DSM 46765
Mycobacterium sp IS-3022
Mycobacterium sp GA-0227b
Mycobacterium rutilum DSM 45405
Mycobacterium flavescens DSM 43991
Mycobacterium malmesburyense WCM 7299
Mycobacterium komanii GPK 1020
Mycobacterium novocastrense DSM 44203
Mycobacterium neumannii CECT 8766
Mycobacterium lehmannii CECT 8763
Mycobacterium agri CCUG 37673
Mycobacterium holsaticum DSM 44478
Mycobacterium pulveris DSM 44222
Mycobacterium elephantis Lipa
Mycobacterium elephantis DSM 44368
Mycobacterium litorale DSM 45785
Mycobacterium doricum CCUG 46352
Mycobacterium sp JLS
Mycobacterium monacense DSM 44395
Mycobacterium sp KMS
Mycobacterium sp MCS
Mycobacterium intermedium DSM 44049
Mycobacterium bourgelatii DSM 45746
Mycobacterium kubicae DSM 44627
Mycobacterium persicum AFPC-000227
Mycobacterium kansasii ATCC 12478
Mycobacterium marinum M
Mycobacterium marinum CCUG 20998
Mycobacterium marinum 1218R
Mycobacterium mantenii DSM 45255
Mycobacterium arosiense ATCC BAA-1401=DSM 45069
Mycobacterium colombiense CECT 3035
Mycobacterium avium subsp paratuberculosis MAP4
Mycobacterium avium 104
Mycobacterium timonense CCUG 56329T
Mycobacterium bouchedurhonense DSM 45439
Mycobacterium parmense DSM 44553
Mycobacterium paraense DSM 46749
Mycobacterium nebraskense AKUC1
Mycobacterium nebraskense DSM 44803
Mycobacterium paraffinicum DSM 44181
Mycobacterium europaeum DSM 45397
Mycobacterium seoulense DSM 44998
Mycobacterium paraseoulense DSM 45000
1
2.1
2.2

### Figure S4c

*selC*

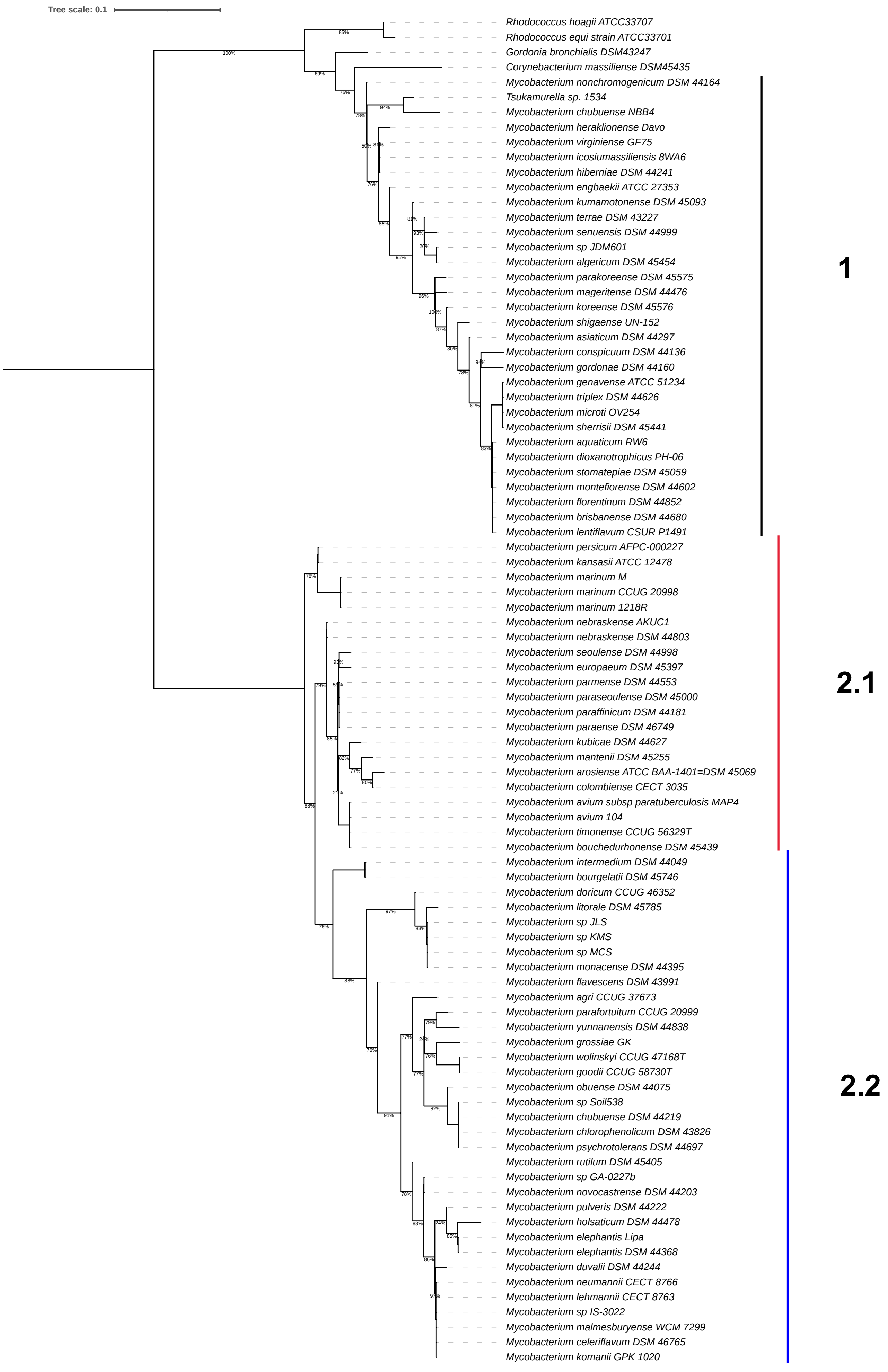

### Figure S4d

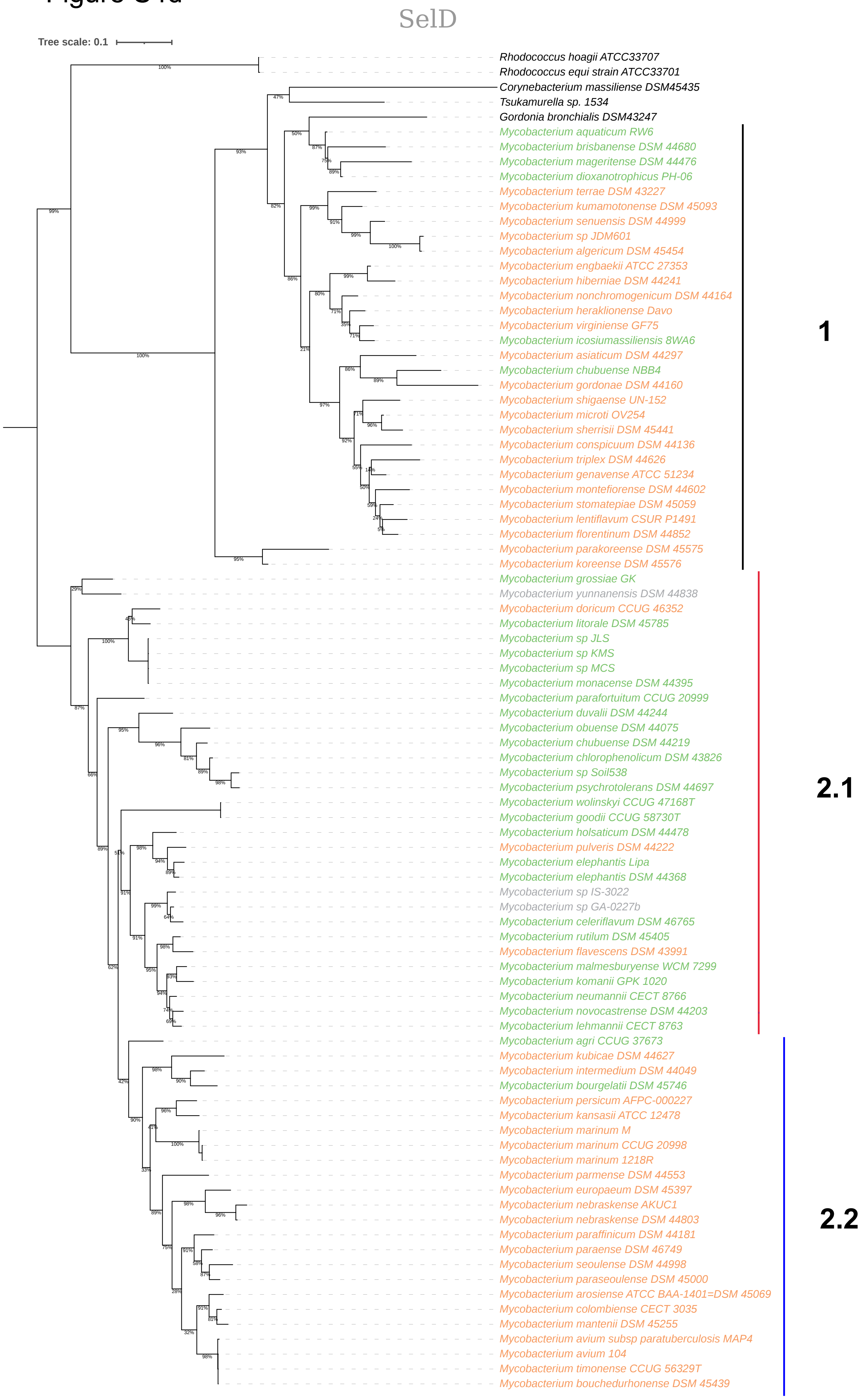

SelD
Tree scale: 0.1
Rhodococcus hoagii ATCC33707
Rhodococcus equi strain ATCC33701
Corynebacterium massiliense DSM45435
Tsukamurella sp. 1534
Gordonia bronchialis DSM43247
Mycobacterium aquaticum RW6
Mycobacterium brisbanense DSM 44680
Mycobacterium mageritense DSM 44476
Mycobacterium dioxanotrophicus PH-06
Mycobacterium terrae DSM 43227
Mycobacterium kumamotonense DSM 45093
Mycobacterium senuensis DSM 44999
Mycobacterium sp JDM601
Mycobacterium algericum DSM 45454
Mycobacterium engbaekii ATCC 27353
Mycobacterium hiberniae DSM 44241
Mycobacterium nonchromogenicum DSM 44164
Mycobacterium heraklionense Davo
Mycobacterium virginiense GF75
Mycobacterium icosiumassiliensis 8WA6
Mycobacterium asiaticum DSM 44297
Mycobacterium chubuense NBB4
Mycobacterium gordonae DSM 44160
Mycobacterium shigaense UN-152
Mycobacterium microti OV254
Mycobacterium sherrisii DSM 45441
Mycobacterium conspicuum DSM 44136
Mycobacterium triplex DSM 44626
Mycobacterium genavense ATCC 51234
Mycobacterium montefiorense DSM 44602
Mycobacterium stomatepiae DSM 45059
Mycobacterium lentiflavum CSUR P1491
Mycobacterium florentinum DSM 44852
Mycobacterium parakoreense DSM 45575
Mycobacterium koreense DSM 45576
Mycobacterium grossiae GK
Mycobacterium yunnanensis DSM 44838
Mycobacterium doricum CCUG 46352
Mycobacterium litorale DSM 45785
Mycobacterium sp JLS
Mycobacterium sp KMS
Mycobacterium sp MCS
Mycobacterium monacense DSM 44395
Mycobacterium parafortuitum CCUG 20999
Mycobacterium duvalii DSM 44244
Mycobacterium obuense DSM 44075
Mycobacterium chubuense DSM 44219
Mycobacterium chlorophenolicum DSM 43826
Mycobacterium sp Soil538
Mycobacterium psychrotolerans DSM 44697
Mycobacterium wolinskyi CCUG 47168T
Mycobacterium goodii CCUG 58730T
Mycobacterium holsaticum DSM 44478
Mycobacterium pulveris DSM 44222
Mycobacterium elephantis Lipa
Mycobacterium elephantis DSM 44368
Mycobacterium sp IS-3022
Mycobacterium sp GA-0227b
Mycobacterium celeriflavum DSM 46765
Mycobacterium rutilum DSM 45405
Mycobacterium flavescens DSM 43991
Mycobacterium malmesburyense WCM 7299
Mycobacterium komanii GPK 1020
Mycobacterium neumannii CECT 8766
Mycobacterium novocastrense DSM 44203
Mycobacterium lehmannii CECT 8763
Mycobacterium agri CCUG 37673
Mycobacterium kubicae DSM 44627
Mycobacterium intermedium DSM 44049
Mycobacterium bourgelatii DSM 45746
Mycobacterium persicum AFPC-000227
Mycobacterium kansasii ATCC 12478
Mycobacterium marinum M
Mycobacterium marinum CCUG 20998
Mycobacterium marinum 1218R
Mycobacterium parmense DSM 44553
Mycobacterium europaeum DSM 45397
Mycobacterium nebraskense AKUC1
Mycobacterium nebraskense DSM 44803
Mycobacterium paraffinicum DSM 44181
Mycobacterium paraense DSM 46749
Mycobacterium seoulense DSM 44998
Mycobacterium paraseoulense DSM 45000
Mycobacterium arosiense ATCC BAA-1401=DSM 45069
Mycobacterium colombiense CECT 3035
Mycobacterium mantenii DSM 45255
Mycobacterium avium subsp paratuberculosis MAP4
Mycobacterium avium 104
Mycobacterium timonense CCUG 56329T
Mycobacterium bouchedurhonense DSM 45439
1
2.1
2.2

### Figure S4e

#### FdhA

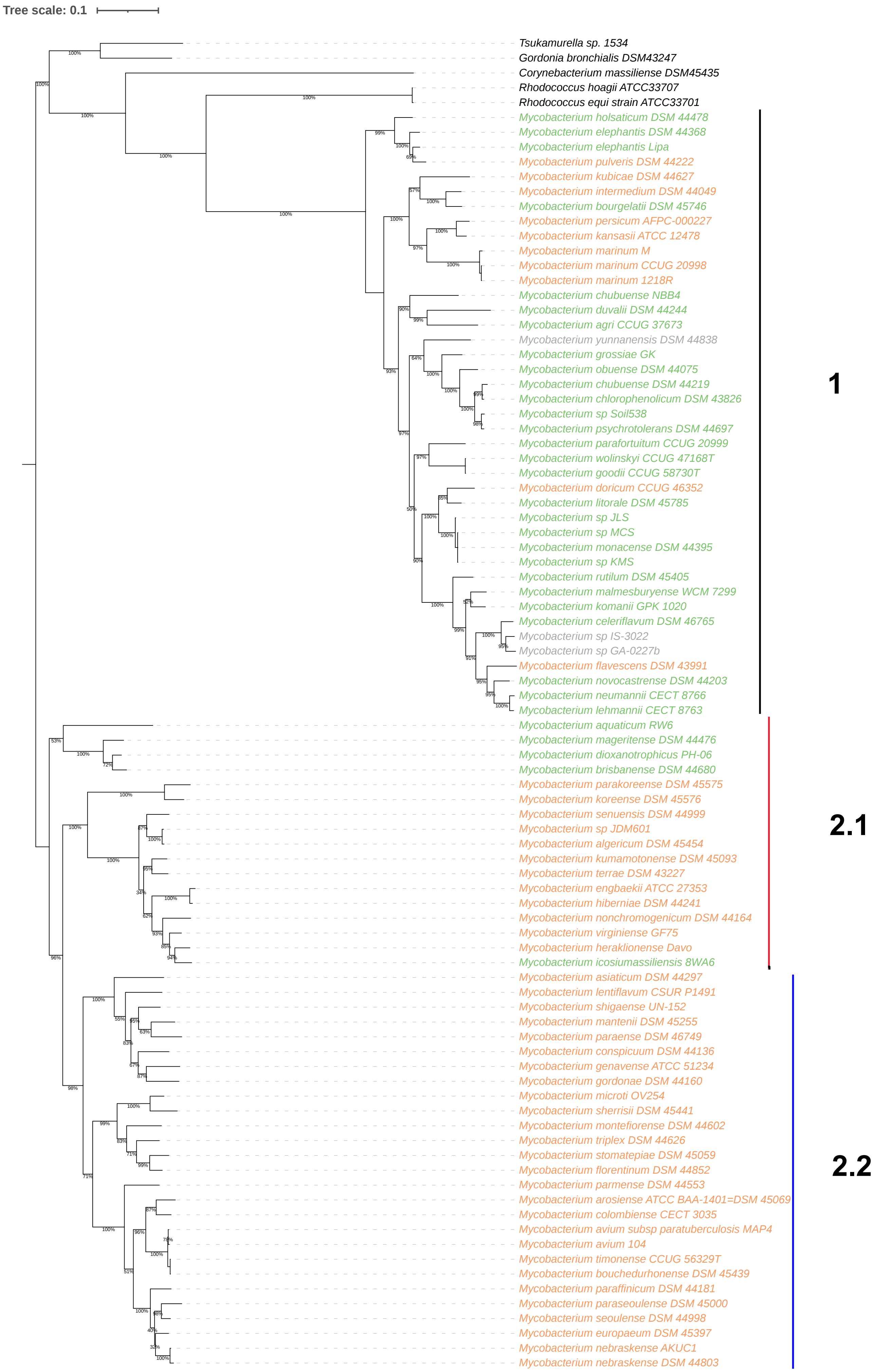

**Figure S5**

Transcript levels of SeC-machinery genes and *sigE* in the *M. marinum* 1218R, 1218S and M strains as indicated.

(a) Distribution of SeC-machinery, *fdhA* and $tRNA^{SeC}$ transcript levels in exponential and stationary cells shown as TPM values (see Materials and Methods).

(b) TPM values for *sigE* mRNA levels in exponential and stationary cells.

(c) Change in SeC-machinery and *fdhA* mRNA levels, and $tRNA^{SeC}$ levels, comparing levels
in exponentially growing cells and stationary cells. The values are expressed in $\log_2$-fold change as indicated. See main text for details and Materials and Methods.

Statistical significance: *p+adj< 0.05; **: p+adj < 0.01; ***: p+adj< 0.001.

### Figure S5

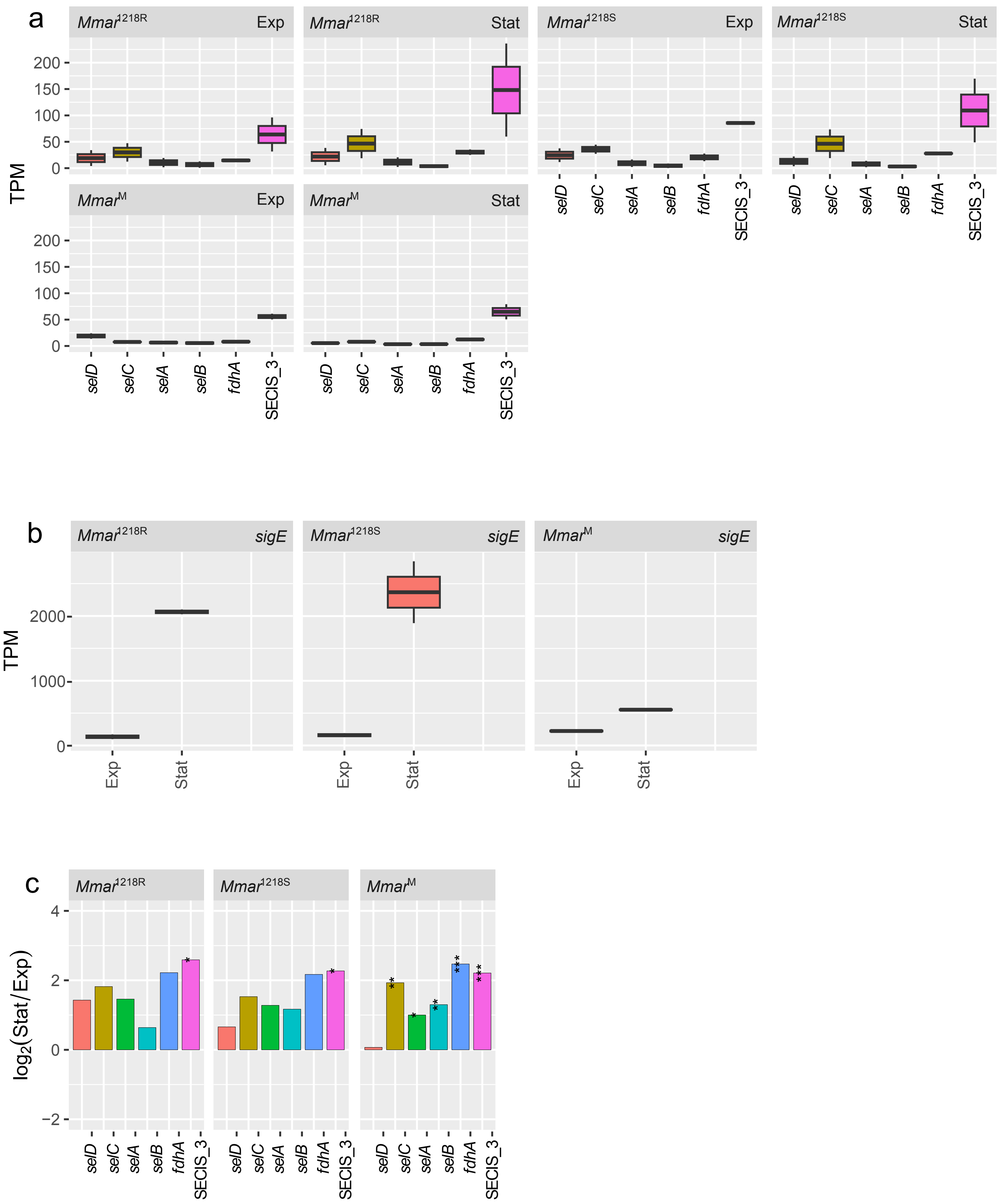

a
Mmar1218R Exp
Mmar1218R Stat
Mmar1218S Exp
Mmar1218S Stat
MmarM Exp
MmarM Stat
TPM
200
150
100
50
0
selD
selC
selA
selB
fdhA
SECIS_3
b
Mmar1218R sigE
Mmar1218S sigE
MmarM sigE
TPM
2000
1000
0
Exp
Stat
c
Mmar1218R
Mmar1218S
MmarM
log2(Stat/Exp)
4
2
0
−2

**Figure S6**

Transcript levels of SeC-machinery genes and *sigE* in the *M. marinum* CCUG strain in response to exposure to different stress conditions – acidic, cold, heat, osmotic and oxidative stress, and exposure to isoniazid, $O_2$-depletion, mitomycin C and starvation as indicated. For details about the conditions, see [37].

(a) Distribution of SeC-machinery, *fdhA* and $tRNA^{SeC}$ transcript levels in exponential and stationary cells shown as TPM values (see Materials and Methods).

(b) TPM values for *sigE* mRNA levels in exponential and stationary cells.

(c) Change in SeC-machinery and *fdhA* mRNA levels, and $tRNA^{SeC}$ levels, comparing levels in exponentially growing cells and stationary cells. The values are expressed in $log_2$-fold change as indicated. See main text for details and Materials and Methods.

Statistical significance: *p+adj< 0.05; **: p+adj < 0.01; ***: p+adj< 0.001.

### Figure S6

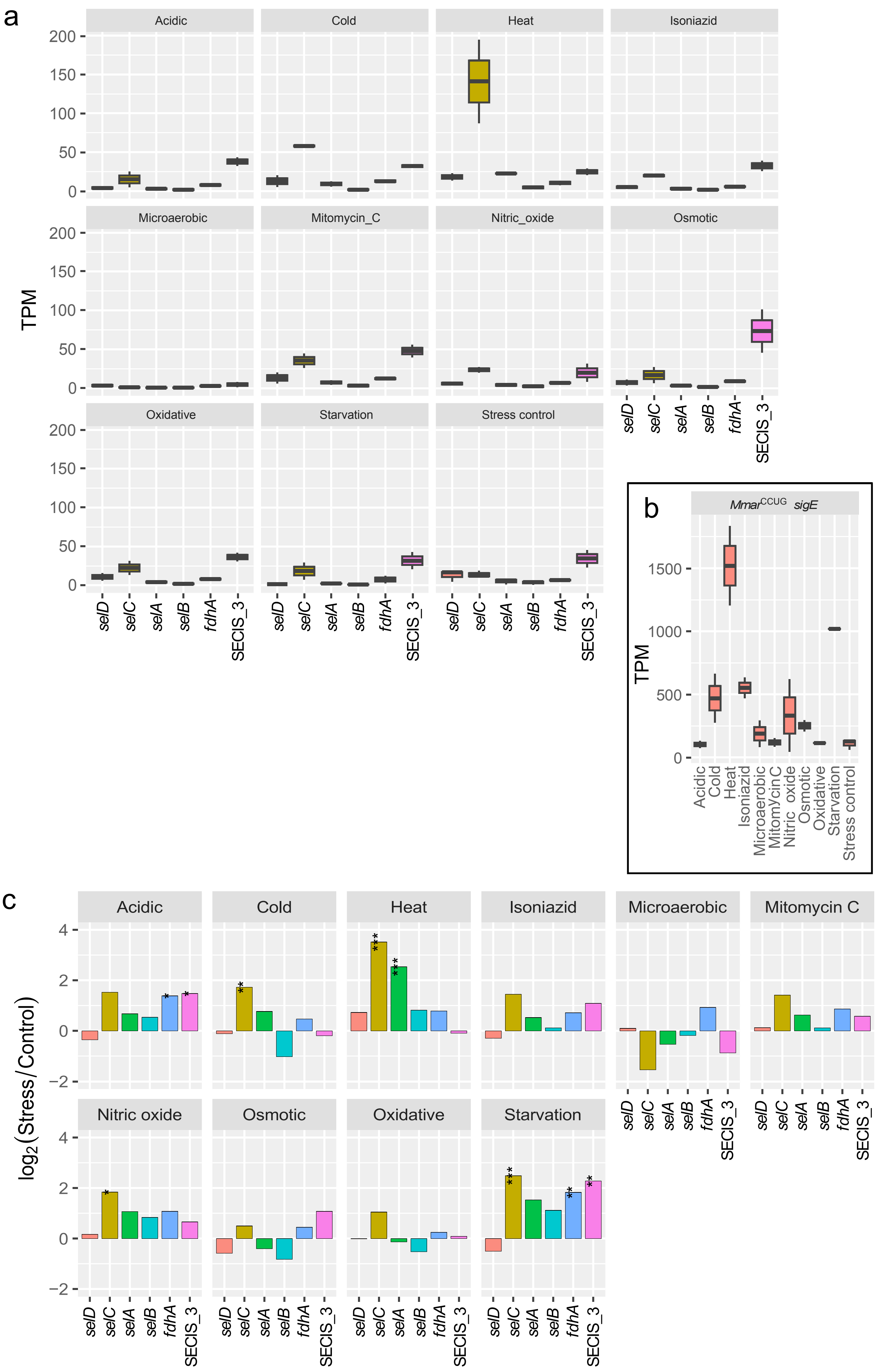
a
Acidic
Cold
Heat
Isoniazid
Microaerobic
Mitomycin_C
Nitric_oxide
Osmotic
Oxidative
Starvation
Stress control
TPM
selD
selC
selA
selB
fdhA
SECIS_3
b
MmarCCUG sigE
TPM
Acidic
Cold
Heat
Isoniazid
Microaerobic
MitomycinC
Nitric_oxide
Osmotic
Oxidative
Starvation
Stress control
c
Acidic
Cold
Heat
Isoniazid
Microaerobic
Mitomycin C
Nitric oxide
Osmotic
Oxidative
Starvation
log2(Stress/Control)

**Figure S7**

Transcript levels of SeC-machinery genes and *sigE* in selected mycobacteria.

(a-c) *Mmon*$^{pD5}$ - 3 days, 6 days, 14 days and 48 days old cells.

(d-f) *Mele* – 2.5 days, 5.5 days, 17 days and 31 days old cells.

(g-i) *Mmfi* – 10 days and 24 weeks old cells, *Mmag* – 1 day and 10 days old cells, as indicated.

(a, d and g) Distribution of SeC-machinery, *fdhA* and tRNA$^{SeC}$ transcript levels in exponential and stationary cells shown as TPM values (see Materials and Methods).

(b, e and h) TPM values for *sigE* mRNA levels in exponential and stationary cells.

(c, f and i) Change in SeC machinery and *fdhA* mRNA levels, and tRNA$^{SeC}$ levels, comparing levels in exponentially growing cells and stationary cells. The *Mmon*$^{pD5}$ and *Mele* transcript levels were relative to the earliest time points, *i.e.* 3 and 2.5 days, respectively. The values are expressed in $\log_2$-fold change as indicated. See main text for details and Materials and Methods.

Statistical significance: *p+adj< 0.05; **: p+adj < 0.01; ***: p+adj< 0.001.

### Figure S7

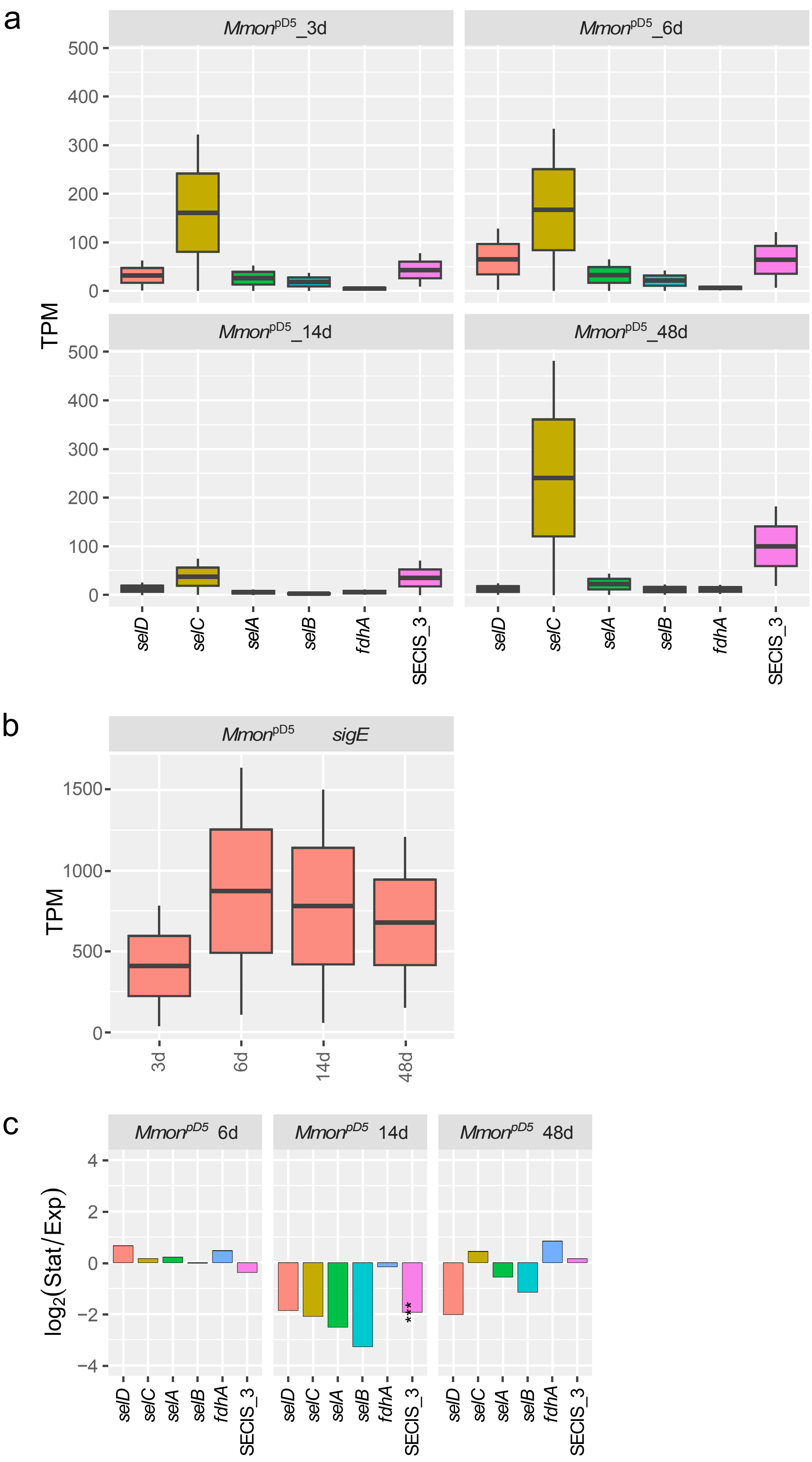
a
Mmon^pD5_3d
Mmon^pD5_6d
Mmon^pD5_14d
Mmon^pD5_48d
TPM
0
100
200
300
400
500
selD
selC
selA
selB
fdhA
SECIS_3
b
Mmon^pD5 sigE
TPM
0
500
1000
1500
3d
6d
14d
48d
c
Mmon^pD5 6d
Mmon^pD5 14d
Mmon^pD5 48d
log2(Stat/Exp)
−4
−2
0
2
4
***

**Figure S8**

Transcript levels of SeC-machinery genes and *sigE* in *Mmar*$^{CCUG}$ and *Mmon*$^{pD5}$ in response to exposure to $O_2$-depletion as a function of time. For details about the conditions [37].

(a) *Mmar*$^{CCUG}$, distribution of SeC-machinery, *fdhA* and tRNA$^{SeC}$ transcript levels in exponential and stationary cells shown as TPM values (see Materials and Methods).

(b) *Mmar*$^{CCUG}$, TPM values for *sigE* mRNA levels in exponential and stationary cells.

(c) *Mmon*$^{pD5}$, distribution of SeC-machinery, *fdhA* and tRNA$^{SeC}$ transcript levels in exponential and stationary cells shown as TPM values (see Materials and Methods).

(d) *Mmon*$^{pD5}$, TPM values for *sigE* mRNA levels in exponential and stationary cells.

Figure S8

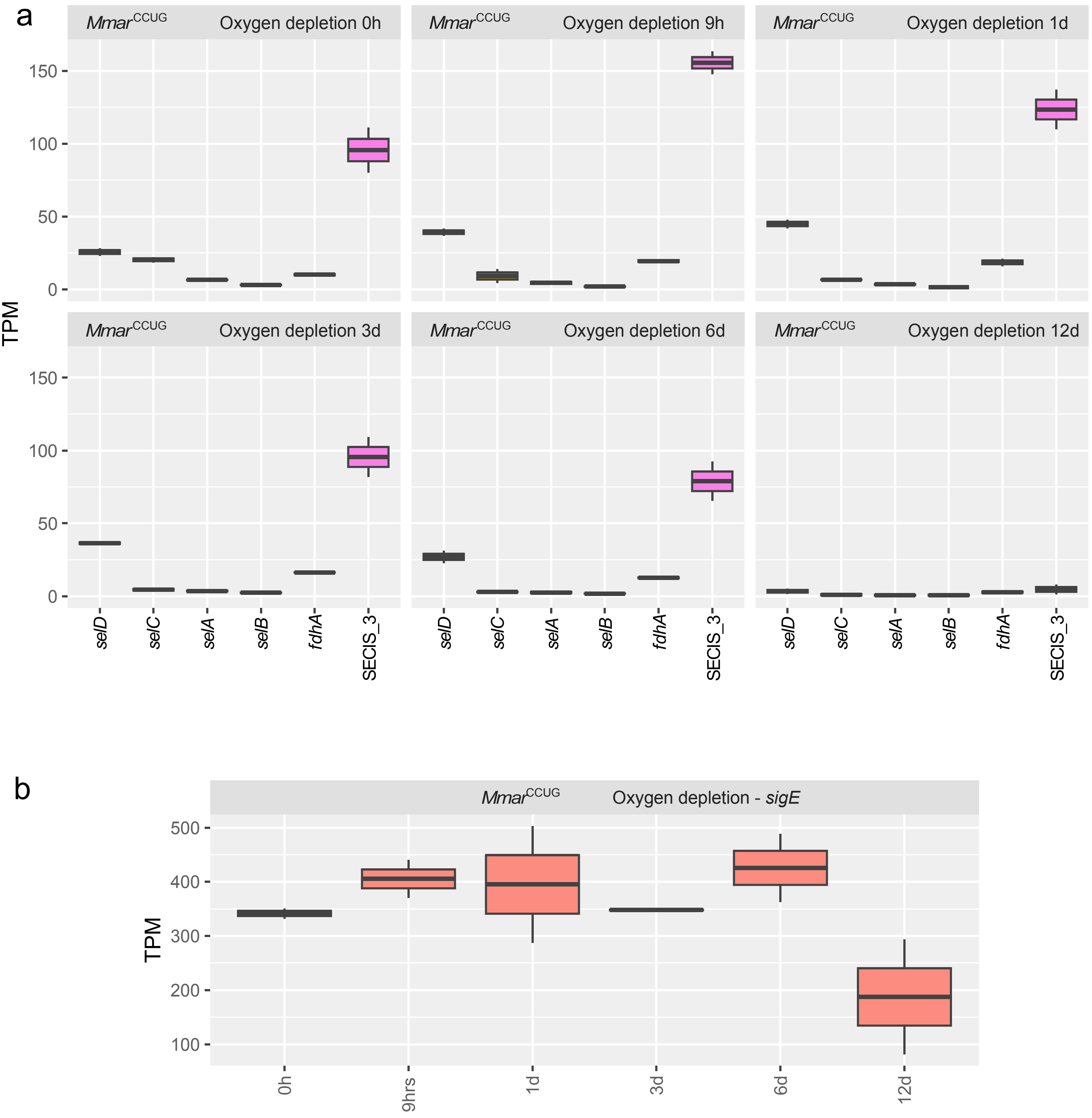
a
Mmar CCUG Oxygen depletion 0h
Mmar CCUG Oxygen depletion 9h
Mmar CCUG Oxygen depletion 1d
Mmar CCUG Oxygen depletion 3d
Mmar CCUG Oxygen depletion 6d
Mmar CCUG Oxygen depletion 12d
TPM
0
50
100
150
selD
selC
selA
selB
fdhA
SECIS_3
b
Mmar CCUG Oxygen depletion - sigE
TPM
100
200
300
400
500
0h
9hrs
1d
3d
6d
12d

Figure S8

c

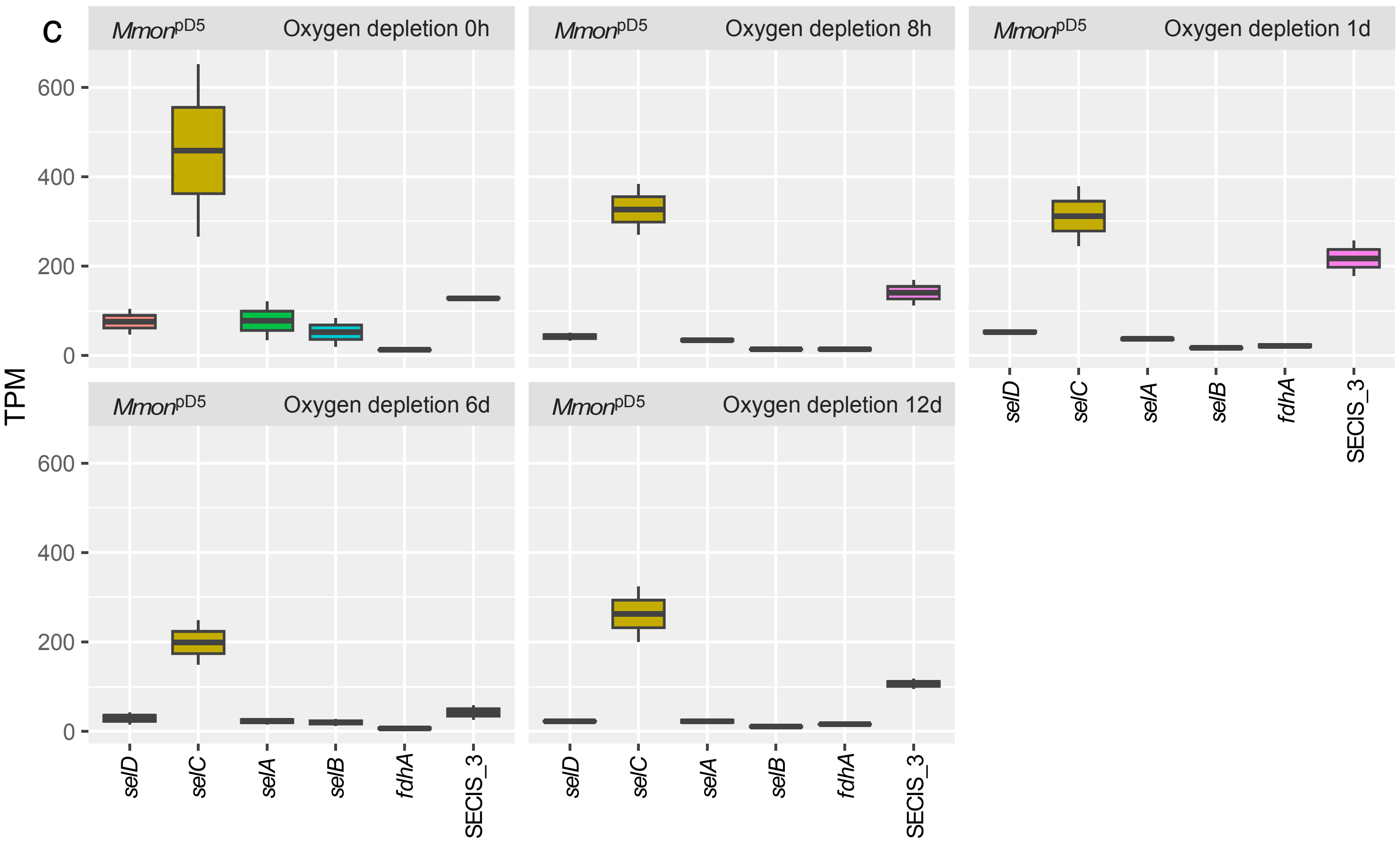

Mmon pD5 Oxygen depletion 0h
Mmon pD5 Oxygen depletion 8h
Mmon pD5 Oxygen depletion 1d
Mmon pD5 Oxygen depletion 6d
Mmon pD5 Oxygen depletion 12d
TPM
600
400
200
0
selD
selC
selA
selB
fdhA
SECIS_3


d

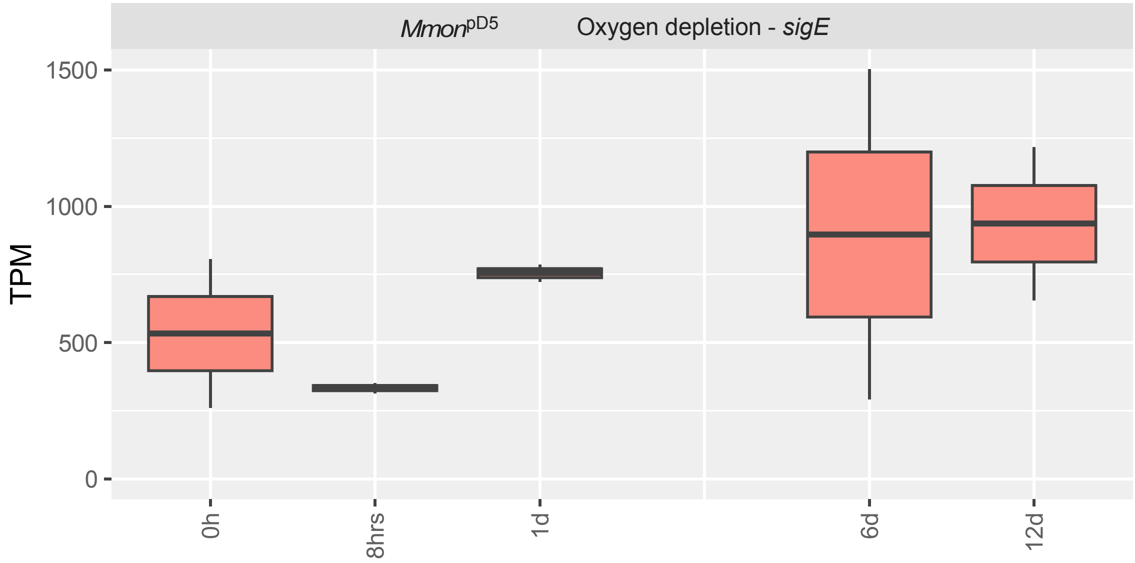

Mmon pD5 Oxygen depletion - sigE
TPM
1500
1000
500
0
0h
8hrs
1d
6d
12d

**Figure S9**

Snapshots of the aligned paired end reads (indicated in grey and red) for *selD*, *selC* (tRNA$^{SeC}$ gene) and *selA* coding regions. Reads and paired end reads that start in one of the genes and end in the other while spanning the region between them are indicated with black arrows. The bottom tracks coloured in blue indicate *selD*, *selC* (tRNA$^{SeC}$ gene) and *selA*. Top panels represent reads in exponential growing cells and bottom panels stationary cells.

(a) *Mmar*$^{CCUG}$

(b) *Mmar*$^{1218R}$

(c) *Mmar*$^{1218S}$

(d) *Mmar*$^{M}$

(e) *Mele*

(f) *Mmfi*

(g) *Mmon*

(h) *Mmag*

### Figure S9

a

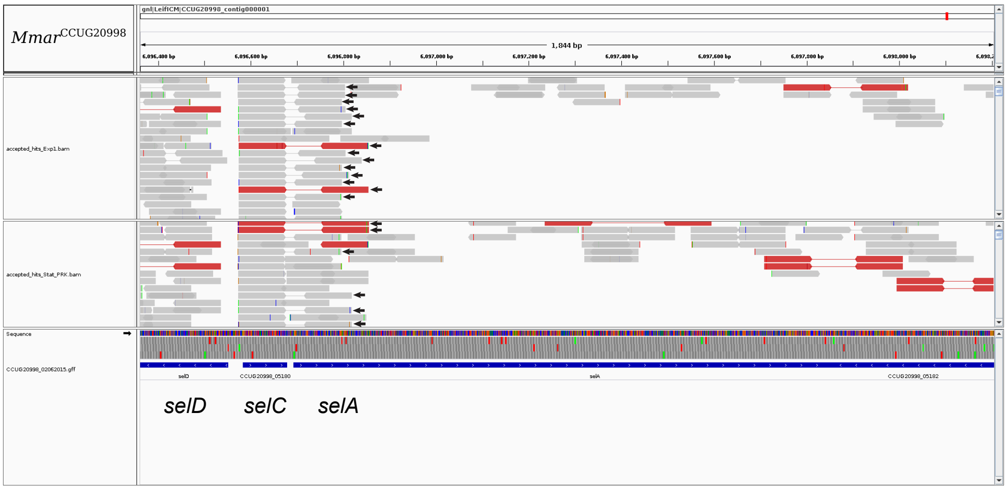


b

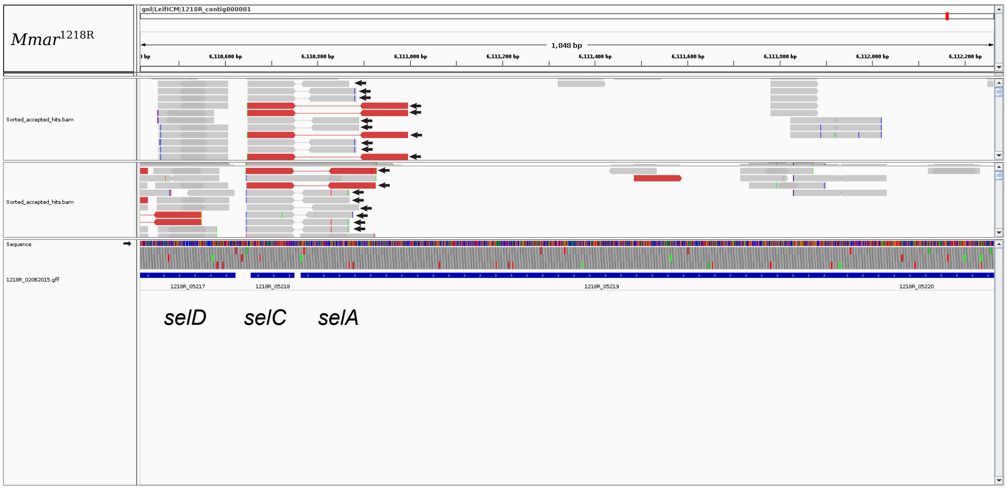


c

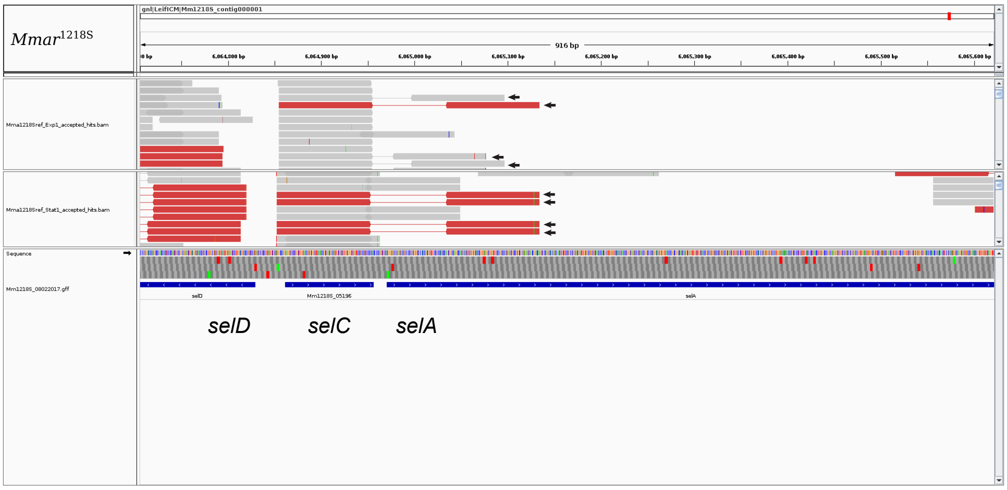


d

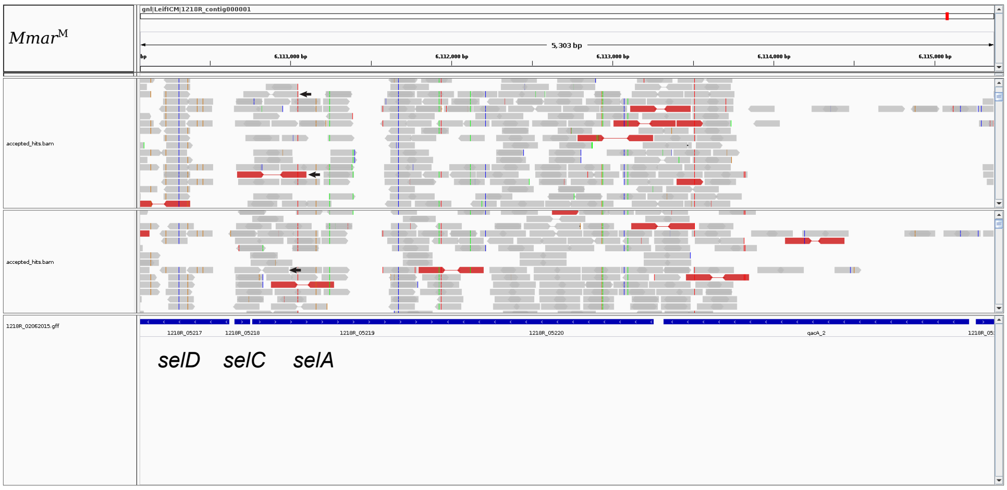

### Figure S9

e

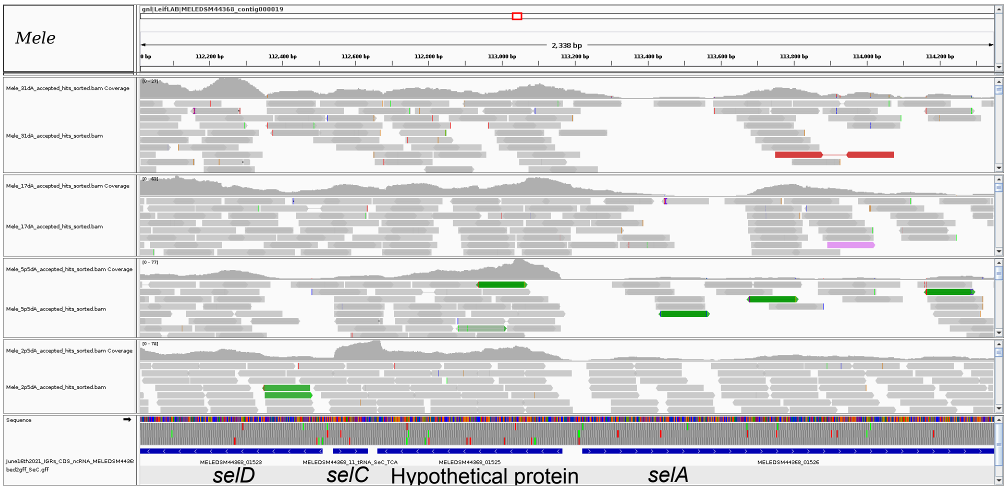


f

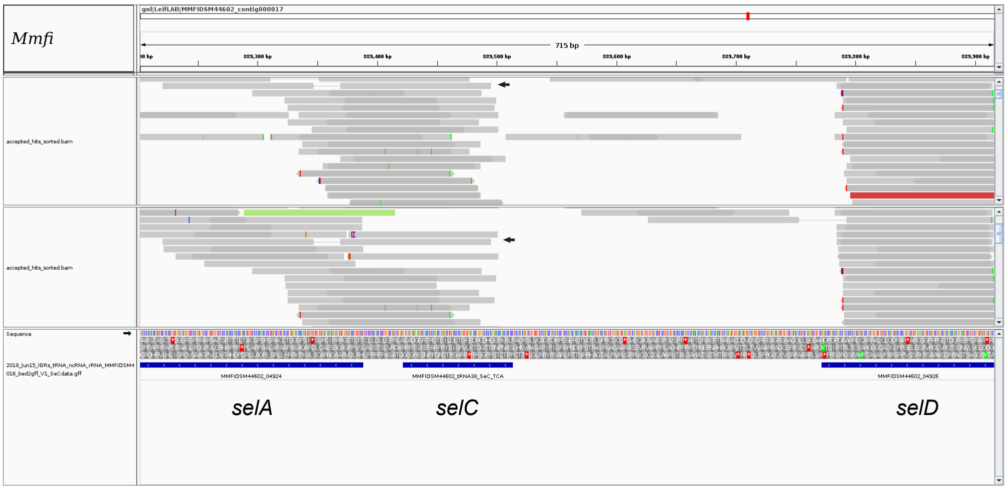


g

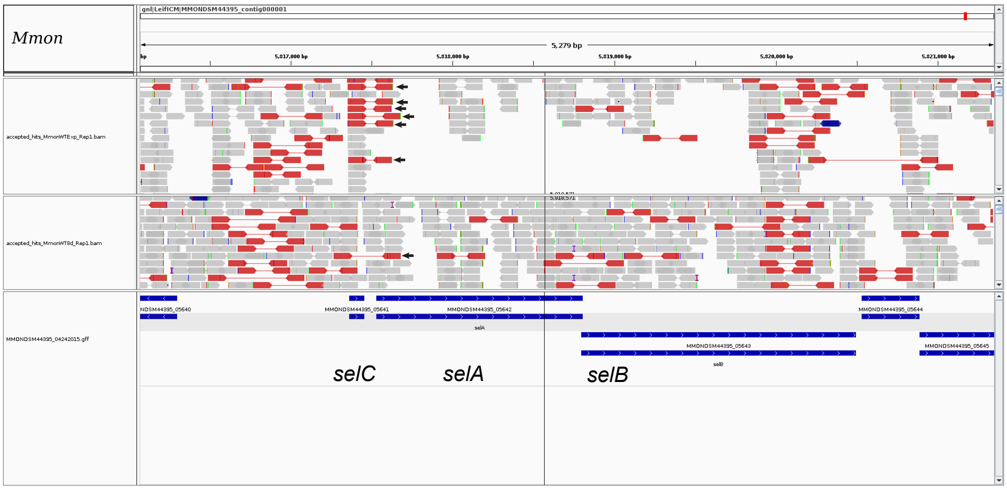


h

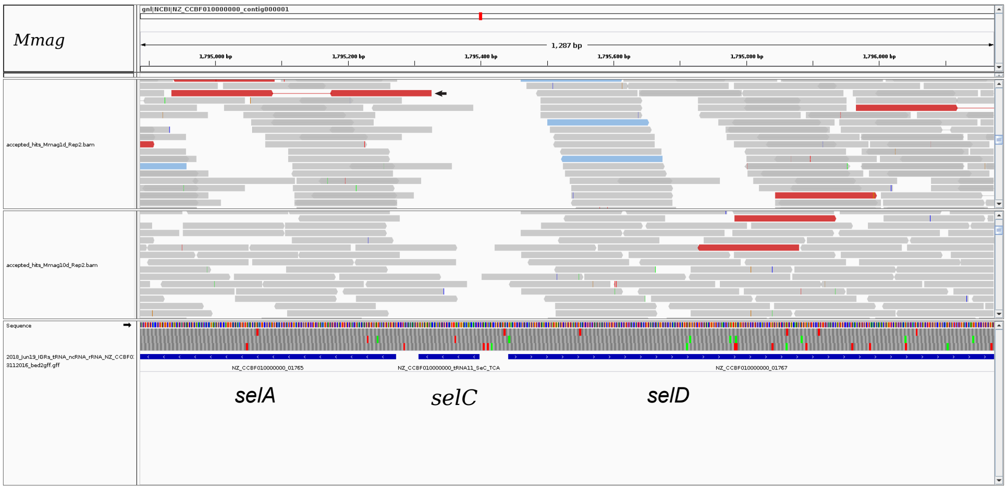
